# NOTCH3 Modulation of Extracellular Matrix, Cytoskeletal Organisation and Metabolic Functions in Human Vascular Smooth Muscle Cells

**DOI:** 10.64898/2026.08.27.746276

**Authors:** Stephen Fitzsimons, Eugene Dillon, Darrell Andrews, Keith J. Murphy, Eoin Brennan, Fanny M. Elahi, Catherine Godson

**Author notes:** **Corresponding Author:** Stephen Fitzsimons. **Data Availability Statement:** Data is available upon reasonable request. Proteomics data is located in PRIDE.

## Abstract

NOTCH3 is a transmembrane receptor highly expressed in vascular mural cells where it contributes to blood vessel formation and homeostasis. NOTCH3 expression declines in the vasculature with aging, and dysregulated NOTCH3 signalling is implicated in pulmonary arterial hypertension, cancer progression and CADASIL (Cerebral Autosomal Dominant Arteriopathy with Subcortical Infarcts and Leukoencephalopathy). RNA-based approaches targeting NOTCH3 are emerging as potential therapeutic strategies, however, the consequences of NOTCH3 suppression in mature vascular smooth muscle cells (VSMCs) remain incompletely understood. Here, we investigated the molecular and functional effects of siRNA-mediated NOTCH3 knockdown in human aortic smooth muscle cells.

Transfection with NOTCH3-targeting siRNA efficiently suppressed NOTCH3 transcript and protein levels. Quantitative proteomics revealed remodelling of extracellular matrix (ECM), cytoskeletal and metabolic pathways, with enrichment of collagen biosynthesis and inhibition of glycolytic signalling. Specifically, NOTCH3 knockdown increased ECM components, including COL3A1, elevated F-actin, and upregulated the actin regulator, CTTN. In parallel, glycolytic capacity was reduced, accompanied by decreased expression of the glycolytic enzyme ENO2. Despite reduced *VEGFA* and alteration in angiogenic signalling proteins, endothelial network formation in co-cultures, as well as VSMC proliferation and migration remained unaffected. Finally, NOTCH3 interactome analysis revealed key collagen and actin-regulating proteins.

These findings identify NOTCH3 as an important regulator of ECM homeostasis, cytoskeletal organisation, and glycolytic metabolism. The preservation of primary cellular functions despite molecular remodelling highlights the adaptive capacity of VSMCs. These findings demonstrate that therapeutic modulation of NOTCH3 may alter vascular cell biology which warrants consideration during development of RNA-based therapeutics for CADASIL and other NOTCH3-associated diseases.

**Graphical Abstract:** **Effects of NOTCH3 knockdown on human aortic smooth muscle cells (HAoSMCs).** HAoSMCs were treated with NOTCH3 siRNA followed by stimulation with PDGF-BB or TGF-β to model VSMC activation. NOTCH3 knockdown induced molecular remodelling characterised by increased ECM-associated pathways, including COL3A1 and collagen biosynthesis, alongside alterations in cytoskeletal proteins, CTNN and F-Actin. Metabolic alterations were observed, including a decrease in ENO2 and a trend towards decreased glycolytic capacity. NOTCH3 knockdown also increased expression of the epigenetic regulator SETD7. Despite reduction in *VEGFA* expression, there was no change in angiogenic endothelial cell network formation when exposed to conditioned media from the HAoSMCs or in direct contact with HAoSMCs lacking NOTCH3. Functionally, migration and proliferation were also unchanged indicating preservation of key VSMC responses. Collectively, NOTCH3 siRNA causes molecular remodelling while maintaining VSMC functional capacity.

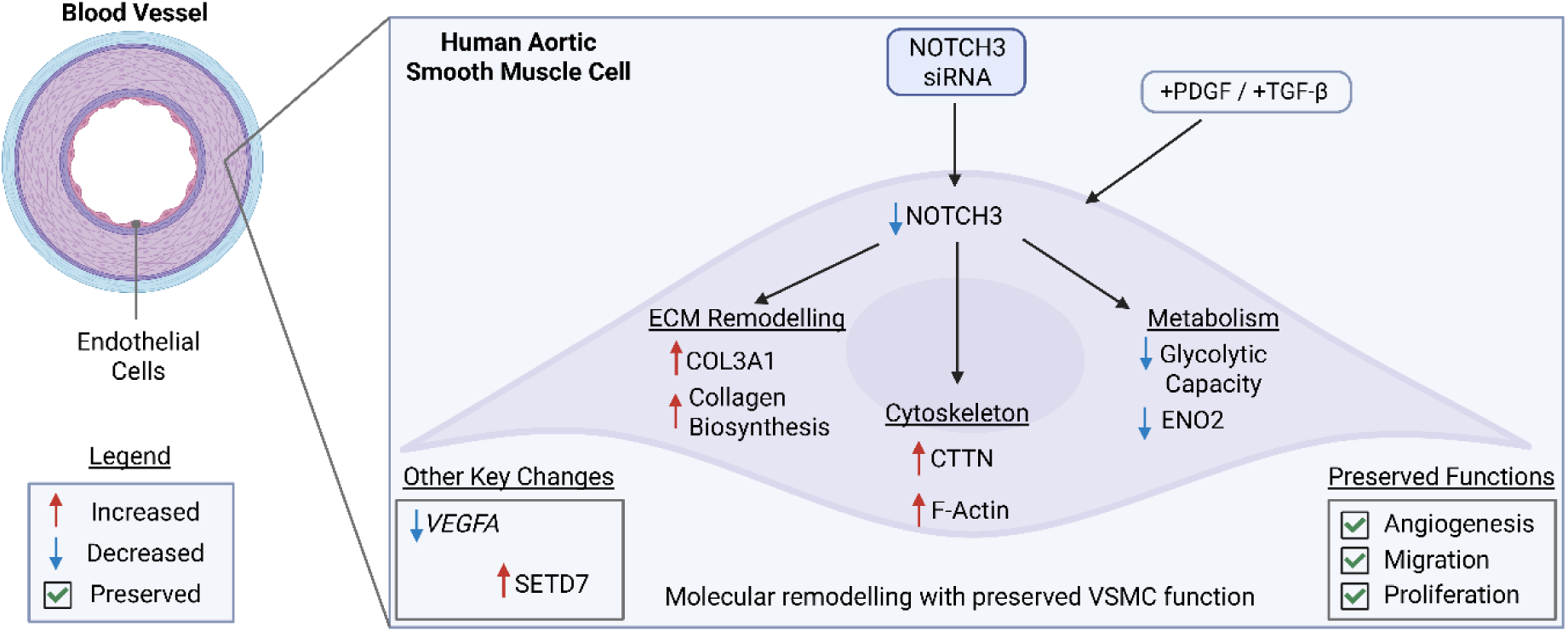

**Key Points:**

- NOTCH3 knockdown increases COL3A1 and total collagen deposition following stimulation with PDGF and TGF-β.
- NOTCH3 knockdown suppresses glycolytic capacity and reduces the glycolytic enzyme ENO2.
- NOTCH3 knockdown increases the abundance of key actin-associated proteins, including cortactin (CTTN) and F-actin.
- The NOTCH3 interactome encompasses structural collagens (COL3A1) and actin-associated regulatory proteins (CTTN).
- NOTCH3 knockdown does not alter angiogenesis, migration, or proliferation under the investigated conditions.

## Introduction

Vascular mural cells, comprising microvessel-associated pericytes and large vessel-associated vascular smooth muscle cells (VSMCs), interact closely with endothelial cells (ECs) to regulate the function, formation, stabilization and remodelling of vasculature [1]. VSMCs exhibit phenotypic plasticity, existing along a dynamic continuum between contractile and synthetic states. These distinct functional states are characterized by specific gene and protein expression profiles reflecting alterations in morphology, proliferation and migration rates. This plasticity enables MCs to maintain vascular homeostasis under physiological conditions while adapting to vascular injury and disease [2]. Following vascular injury, VSMCs switch to a ‘synthetic’, highly proliferative state that promotes ECM deposition and tissue repair before reacquiring a contractile phenotype during the restoration of vessel homeostasis [3]. Transitions between these phenotypic states are regulated by local microenvironmental cues, intrinsic genetic programming and epigenetic mechanisms [4–6].

NOTCH3 is a highly conserved, single pass, transmembrane signalling receptor that is predominantly expressed in VSMCs and pericytes where it plays a central role in vascular development and homeostasis [7, 8]. The receptor contains a large extracellular domain (ECD) composed of Epidermal Growth Factor-like repeats (EGFr) that interact with membrane bound ligands of the Jagged (JAG1 and JAG2) and Delta-like (DLL1, DLL3 and DLL4) families expressed on neighbouring cells. Ligand binding initiates sequential proteolytic cleavage, releasing the NOTCH3 intracellular domain (ICD), which translocates to the nucleus mediating gene expression [9, 10].

NOTCH signalling co-ordinates vessel development through VSMC-EC and VSMC-VSMC cell-cell communication [11, 12]. During angiogenesis, EC-expressed JAG1 initiates heterotypic, EC-VSMC, signalling by activating NOTCH3 in co-cultured mural cells [11]. Crucially, this primary event activates a feed-forward mechanism of lateral induction wherein mural cells upregulate both NOTCH3 and JAG1, enabling homotypic, VSMC-VSMC signalling that propagates and consolidates the differentiated, mature VSMC arterial phenotype along the vessel wall [13–15]. Consequently, NOTCH3 is a regulator of arterial maturation, mural cell differentiation and vascular remodelling.

Progressive loss of NOTCH3 and JAG1 is reported in the aging murine cerebrovasculature, while NOTCH3 expression similarly declines with age in the human cerebrovasculature [16]. Age-related decline in NOTCH3 signalling leads to structural deterioration of the vessel wall, characterised by progressive disorganization, phenotypic dedifferentiation, and detachment of VSMCs. Furthermore, loss of NOTCH3 in VSMCs results in a downregulation in contractile and extracellular matrix (ECM)-associated genes [16].

Dysregulation of NOTCH3 signalling has been implicated in several pathological conditions including pulmonary arterial hypertension (PAH), CADASIL and cancer. In PAH, NOTCH3 signalling contributes to pathological vascular remodelling with increased NOTCH3 expression observed in pulmonary artery SMCs from both patients and experimental models of pulmonary hypertension [17]. Pulmonary NOTCH3 protein levels correlate with disease severity, while genetic deletion of Notch3 or pharmacological inhibition of NOTCH activation attenuates hypoxia-induced pulmonary hypertension in mice [17]. Furthermore, serum NOTCH3 extracellular domain (NOTCH3-ECD) levels have demonstrated potential as a biomarker of idiopathic PAH, correlating with disease severity and progression [18].

Beyond vascular disease, aberrant NOTCH3 activation has also been implicated in tumour progression where it can influence cell differentiation, tissue commitment, cell proliferation and invasion (reviewed in [19]). For example, NOTCH3 knockdown selectively reduced survival of metastatic-derived head and neck squamous cell carcinoma cell lines and limited tumour growth *in vivo* [20]. Additionally, inhibition of NOTCH signalling in tumour-derived mesenchymal cells altered ECM production, reducing expression of collagen (COL1A1) and connective tissue growth factor (CTGF), while impairing invasive capacity [21]. Finally NOTCH3 inhibition has been shown to reduce tumour cell colony formation [22]. Collectively, there is a context dependent role of NOTCH3 signalling in regulating cellular phenotype and tissue remodelling across diverse disease settings. In certain cancers, pathological NOTCH3 activation contributes to tumour progression and resistance to therapy, suggesting that therapeutic inhibition of NOTCH3 may provide clinical benefit.

As a monogenic disorder caused by pathogenic variants in NOTCH3, CADASIL provides a unique model for investigating the physiological functions of NOTCH3 signalling. The majority of CADASIL-causing mutation occur within the EGFr of the NOTCH3 ECD, disrupting the conserved cysteine residue pattern and generating an unpaired cysteine that promotes receptor misfolding and aberrant binding to other proteins causing aggregation [23]. The resulting cerebral small vessel pathology is characterized by accumulation of the NOTCH3 ECD and granular osmiophilic deposits, progressive VSMC degeneration, altered distribution of VSMCs, excessive ECM deposition and vascular fibrosis, ultimately leading to vessel wall thickening, impaired cerebrovascular reactivity, and reduced cerebral blood flow [24–28]. The clinical severity of CADASIL is influenced by the location of the cysteine altering variant withing the EGFr domain [29]. Clinically, CADASIL presents with fatigue, migraine with aura, recurrent subcortical ischaemic strokes, mood disturbances, apathy, cognitive impairment, and progressive vascular dementia [30–32].

A central question in CADASIL pathobiology is whether the disease results primarily from impaired NOTCH3 signalling (loss of function), a toxic gain-or neomorphic function arising from mutant NOTCH3 accumulation [33], or a combination of both mechanisms [34]. The extracellular accumulation of the NOTCH3 ECD and the formation of GOMs are pathological hallmarks and support a gain of function mechanism, whereby mutant NOTCH3 forms extracellular aggregates that sequester multiple matrix and vascular proteins [33, 35, 36], thereby disrupting vessel wall homeostasis. Furthermore, most pathogenic NOTCH3 variants occur within EGFr that exhibit relatively low evolutionary sequence conservation, suggesting that many mutations may preserve the core signalling function of the receptor (ligand binding and cleavage), while promoting protein misfolding and aggregation [37]. However, several studies have reported that CADASIL-associated mutations also impair normal NOTCH3 signalling [38–40], potentially compromising VSMC differentiation, maintenance, and function. Additionally, transendocytosis of mutant NOTCH3 has been reported to be impaired in some CADASIL-associated mutations, thereby prolonging its half-life relative to wild-type NOTCH3, which may alter NOTCH3 ICD generation and downstream transcriptional signalling [41, 42]. The relative contribution of toxic protein accumulation and impaired canonical NOTCH3 signalling to disease pathogenesis therefore remains unresolved. Defining the physiological functions of endogenous NOTCH3 is therefore essential for distinguishing primary loss-of-function effects from secondary consequences of mutant protein accumulation.

Experimental manipulation of NOTCH3 further supports its role in regulating VSMC phenotype and function. In a CADASIL model, iPSC-derived mural cells carrying a NOTCH3 mutation exhibited impaired endothelial tubule network formation. siRNA-mediated knockdown of NOTCH3 restored endothelial tubule network formation and rescued VEGF secretion, demonstrating that reducing mutant NOTCH3 can partially reverse vascular dysfunction [43]. In addition, siRNA-mediated knockdown of NOTCH3 in healthy VSMCs attenuated the EC-induced expression of contractile markers, including *CNN1, ACTA2, TAGLN*, demonstrating that endogenous NOTCH3 signalling mediates EC-VSMC crosstalk to promote VSMC differentiation [11]. Notably, short hairpin RNA-mediated knockdown of *NOTCH3* disrupted actin filament organization in healthy VSMCs, recapitulating the cytoskeletal abnormalities observed in VSMCs derived from patients with CADASIL [27]. In contrast, lentiviral overexpression of the Notch3 intracellular domain in hPSC-derived neural crest cells promoted differentiation into PDGFRβ-positive brain mural cells, accompanied by enhanced ECM production, including fibronectin, and the acquisition of functional ATP-sensitive potassium channels, which contributes to regulation of cerebral vascular tone [44]. Collectively, these studies demonstrate that NOTCH3 is a critical regulator of mural cell differentiation and structural integrity, though its downstream pathways in mature human VSMCs remain incompletely defined.

*In vivo* models have further demonstrated the importance of NOTCH3 for vascular homeostasis. Homozygous *notch3* mutant zebrafish exhibit vasculopathy, oligodendrocyte abnormalities and myelin defects [45] while morpholino-mediated depletion of notch3 reduces the population of *pdgfrb*^high^ mural cells [46]. Similarly, *Notch3* knockout mice display arterial defects, impaired cerebral blood flow reactivity [7], loss of arterial VSMCs through apoptosis, disruptions in blood-brain barrier integrity and fibrin deposition within the retinal vasculature [47]. Collectively, these studies demonstrate that endogenous NOTCH3 is essential for developmental processes, vascular integrity and mural cell homeostasis. In contrast, in a murine CADASIL model, generated through overexpression of the Notch3 R169C mutation on a wild-type background, elimination of one copy of wild-type Notch3 reduced arterial pathology and Notch3 ECD accumulation [33], supporting the therapeutic potential of NOTCH3-lowering strategies. Despite these studies, the molecular and functional consequences of siRNA-mediated knockdown on mature, healthy human VSMCs remain poorly understood, as most evidence to date has focused on developmental processes or disease-associated models.

Several potential therapeutic strategies for CADASIL are currently in preclinical development (reviewed in [48]) including skipping of specific pathogenic NOTCH3 exons using antisense oligonucleotides [49]. Gene silencing of NOTCH3 has emerged as a potential therapeutic approach for both CADASIL and PAH, however, the molecular and cell autonomous role of endogenous NOTCH3 in mature VSMCs has not been fully characterised. Defining the contribution of NOTCH3 to vascular SMC phenotype and function is therefore essential for predicting the efficacy and potential adverse effects of NOTCH3-targeted therapies, particularly as compensatory mechanisms and functional redundancy within the NOTCH signalling pathway may modify cellular responses to gene silencing. Here, we demonstrate that siRNA-mediated knockdown of NOTCH3 in healthy human aortic smooth muscle cels (HAoSMCs) increases collagen, in particular, blood vessel-associated COL3A1, decreases VEGF expression, alters actin cytoskeletal organisation, and impairs aspects of glycolysis. These findings identify NOTCH3 as a key regulator of VSMC homeostasis through coordinated control of ECM remodelling, cytoskeletal organisation and cellular metabolism.

## Materials and Methods

### Cell Culture

Human aortic smooth muscle cells (HAoSMCs) (PromoCell, Cat no. C-12533) were cultured in complete SmBM consisting of SmBM® Basal Medium supplemented with 5% fetal bovine serum (FBS), insulin, human Fibroblast Growth Factor (hFGF), and human Epidermal Growth Factor (hEGF) provided in the SmGM®-2 Smooth Muscle Cell Growth Medium -2 BulletKit® (Lonza, Cat. no. CC-3182) and incubated at 37°C in 5% CO_2_. Human microvascular endothelial cells (HMEC-1) (Cytion, Cat. no. 304064) were cultured in MCDB 131 Medium, no glutamine-500 mL (Cat. no. 10372019) supplemented with FBS, Penicillin-Streptomycin, L-glutamine, hEGF and hydrocortisone.

### siRNA-Mediated Knockdown of NOTCH3

NOTCH3 siRNA-1 and the siRNA Negative Control (Neg Ctrl siRNA) (MedChem Express, Cat. no. HY-150150) from the NOTCH3 Human Pre-designed siRNA Set A (MedChem Express, Cat. no. HY-RS09451) were used for all transfection experiments. HAoSMCs were transfected with NOTCH3 siRNA (20 nM) or Neg Ctrl siRNA (20 nM) using Lipofectamine™ RNAiMAX Transfection Reagent (ThemoFisher Scientific, Cat. no. 13778030) (final concentration of 0.2%), diluted in Opti-MEM I Reduced Serum Medium, no phenol red (ThermoFisher Scientific, Cat No. 31985062) (20%) and SmGM®-2 Smooth Muscle Cell Growth Medium (80%). Untreated HAoSMCs, cultured without lipofectamine, siRNA or Opti-MEM medium were included as a control. Cells were transfected for 6 hours (h) and then rested in SmBM overnight. HAoSMCs were then stimulated for 48 h with PDGF-BB (PDGF) (20 ng/mL) (MedChemExpress, Cat. no. HY-P7055) and TGF beta I (TGF-β) (10 ng/mL) (MedChemExpress, Cat. no. HY-P70543) using PBS with 0.1% BSA as the vehicle control (CTRL).

### Gene Expression Analysis: qRT-PCR

RNA was isolated using the E.Z.N.A. Total RNA Kit I (Omega Biotek, Cat. no. R6834-02) and reverse transcribed using the High-Capacity cDNA Reverse Transcription Kit (Applied BioSystems, Cat. no. 4368814). Quantitative real-time PCR analysis was performed using the QuantStudio™ 5 Real-Time PCR System (ThermoFisher). Results were analyzed via the 2(-Delta Delta C(T)) method [50] using 18S rRNA as the endogenous control. Results were graphed as fold over control (F.O.C.). TaqMan Real-Time PCR Assays used included *COL3A1*, *COL4A1*, *COL6A3*, *CTTN, ENO1*, *IL6, LTBP1*, *NFKB1*, *SETD7*, *SMTN*, *TAGLN* and *VEGFA* while SYBR assays were used for *COL1A1* (see **Supplemental Table 1** for catalogue numbers).

### Western Blotting

Protein was quantified using a Bradford assay (BioRad) according to the manufacturer’s instructions. Western blotting was performed as previously described [51]. In brief, sodium dodecyl sulfate-polyacrylamide gel electrophoresis was used to separated proteins in a polyacrylamide resolving gel (8-10%). A transfer cassette was used to transfer gels to a nitrocellulose membrane. Membranes were blocked for 1 h in skim milk (5%) followed by incubation in primary antibody at 4°C overnight. Secondary antibodies were applied for 1 h at room temperature (RT). Antibody and dilutions and catalogue numbers are available in **Supplemental Table 2**. Membranes were exposed to WesternBright ECL (Advansta, Cat no. K-12045-D20) and developed on a Vilber Fusion Fx Imaging System. β-actin was used as the loading control. Densitometry analysis was performed using ImageJ. Raw blots of all independent experiments are included in **Supplemental Figures 1-3**.

### MTT assay

MTT assays were performed using the *In Vitro* Toxicology Assay Kit, MTT based (Sigma, Cat. no. TOX1-1KT) as per manufacturer’s instructions to assess cell viability through analysis of metabolic activity. Mitomycin-C (MMC) (20 µg/mL) (Fisher BioReagents™, Cat. no. 10182953) was included as the positive control. Plates were quantified spectrophotometrically at 570nm using a CLARIOStar Plate reader (BMG Labtech). Results were normalised to the unstimulated, negative control siRNA.

### Proliferation Assay

HAoSMCs were seeded in a 96-well plate at 1.5×10^3^ cells/well in a volume of 150 µL of complete SmBM. Knockdown of NOTCH3 was performed as outlined previously. Mitomycin C (20 µg/mL) (Fisher BioReagents™, Cat. no. 10182953) and Rotenone (0.5 µM) (Merck, Cat. no. CRM38703) were used as negative controls. Live cell imaging was performed using the Incucyte (Sartorius) for 48 h with images obtained every 4 h.

### Proteomics

The iST kit (Preomics, Cat. no. P.O.00001) was used as per manufacturer’s instructions to isolate peptides from cell samples. Peptides were analysed using a timsTOF mass spectrometer (Bruker). The mass spectrometry proteomics data have been deposited to the ProteomeXchange Consortium via the PRIDE [52] partner repository with the dataset identifier PXD080845 (to be made publicly accessible upon publication).

### Microscopy Analysis

Cells were seeded at 1.5×10^3^ cells/well on PhenoPlate 96-well microplates (formerly CellCarrier Ultra Plates, Perkin Elmer) and following transfection with NOTCH3 siRNA and Neg Ctrl siRNA were stimulated for 48 h with PDGF (20 ng/mL) and TGF-β (10 ng/mL). Cells were stained with MitoTracker Red CMXROS (50 nM) (ThermoFisher Scientific, Cat. no. M7512) for the final hour. Cells were fixed in 4% paraformaldehyde (PFA) and then F-actin was stained using Alex Fluor 488 phalloidin (ThermoFisher Scientific, Cat. no. A12379) for 20 min followed by Hoechst 33342 (ThermoFisher Scientific, Cat. no. H3570) for 10 min. Cells were imaged at 63X using the high content Opera Phenix Automated Screening Microscope (Perkin Elmer). 20 random fields of view (FOV) were obtained per well (60 FOV/well). Three independent experiments were performed with 3 technical replicates per condition. Harmony Imaging Software was used to analyse the images to quantify F-actin and MitoTracker intensity.

### Metabolic Flux Analysis

HAoSMCs were transfected with NOTCH3 siRNA and Neg Ctrl siRNA on 12 well plates for 6 h. Following a rest period of 48 h cells were seeded at 1×10^4^ cells/well in a 96-well Seahorse XF Prom M Cell Culture Microplate. The Agilent Seahorse XF Cell Mito Stress Test was performed using Oligomycin (Oligo) (Sigma, Cat. no. 04876-5mg), Carbonyl cyanide 4-(trifluoromethoxy)phenylhydrazone (FCCP) (MedChemExpress, Cat. no. HY-100410), Rotenone (Merck, Cat. no. CRM38703) and Antimycin (Sigma, Cat. no. A8674-25mg) with a final addition of 2-Deoxy-D-glucose (2-DG, Cat. no. F024330). Three independent experiments were performed with 8-9 technical replicates per condition. Results were analysed using the Seahorse FXPro Analyser and normalised to cell confluence as assessed on Incucyte.

### Total Collagen Analysis

Following treatment of NOTCH3 siRNA and Neg Ctrl siRNA for 6 h as previously described, cells were cultured with PDGF (20ng/mL) and TGF-β (10ng/mL) for 4 days on 96-well plates using PBS with 0.1% BSA as the control (CTRL). Cells were fixed in 4% PFA and washed 3 times with PBS followed by a 1 h incubation with PicoSirius Red Solution (Abcam, Cat. no AB246832) using 100 µL/well and gently shaking. Following removal of the dye, samples were washed 4 times with 0.01N HCL (100 µL/well). Cells were imaged on the Incuyte using phase contrast and the Red Channel (∼585 nm (Ex)/ ∼635 nm(Em)) set to 400 ms acquisition. The PicoSirius Red intensity was quantified relative to the percentage confluence captured by phase contrast. Values were normalised to the percentage confluence.

### Migration Assay/Scratch Wound Assay

HAoSMCs were seeded in an Incucyte Imagelock Plate (96-well) at 3.5×10^3^ cells/well. Cells were transfected for 6 h and then rest as previously described. Cells were treated with Mitomycin C (20 µg/mL) (Fisher BioReagents, Cat. no. 10182953) for 2 h to inhibit cell proliferation. A uniform 700-800 µm scratch was made in each well using the WoundMaker. Cells were cultured in SmBM and PDGF (20 ng/mL) or TGF-β (10 ng/mL) stimulations were included alongside the vehicle CTRL. Live cell imaging was performed using phase contrast microscopy on the Incucyte (Sartorius) for 48 h with images obtained every 4 h. The percentage relative wound density was calculated using the Incucyte software.

### Angiogenesis Assay

Geltrex LDEV-Free Reduced Growth Factor Basement Membrane Matrix (ThermoFisher Scientific, Cat no. 12053569) (35 µL) was applied to a 96 well plate 1 h before seeding cells. HMEC-1 cells were seeded at 1×10^4^ cells/well with and without siRNA transfected HAoSMCs at 0.5×10^4^ cells/well in MCDB 131 Medium (150 µL). For experiments investigating the effects of the HAoSMC supernatant (SN), 100 µL of SN from transfected HAoSMCs cultured for 48 h was added to seeded HMEC-1 cells. Live cell imaging was performed using the Incucyte (Sartorius) for 20 h with images obtained every 1 h. Images from all of the time points were pseudocoloured green using Image J and were analysed by AngioTool2.0 Software to quantify the network formed.

### Immunoprecipitation-Coupled Western Blot and Mass Spectrometry

HAoSMCs were cultured in 10 cm plates at a concentration of 1×10^4^ cells per cm^2^. Cells were cultured for 48 h and the harvested using IP Lysis Buffer (ThermoFisher, Cat no. 87787). Protein was quantified using a Bradford assay (BioRad). Dynabeads Protein G (ThermoFisher Scientific, Cat no. 10003D) were washed twice with lysis buffer. To bind the antibody to the beads, NOTCH3 polyclonal antibody (8 µg total) (Abcam, Cat. no. ab23426) was incubated with 30 µL of pre-washed Dynbeads Protein G for 18 h at 4°C with end-to-end rotation. A rabbit IgG isotype control monoclonal antibody (Brennan & Co, Cell Signalling, Cat. no. 3900S) was used in parallel as the control. Following PBS washes, NOTCH3 antibody-coated beads (30 µL) were mixed with 300 µg of protein and incubated for 18 h at 4°C with end-to-end rotation. Beads were washed 3 times using the magnetic separation rack. The sample was divided for elution for:

1. ***Western Blot***: 33% of sample was eluted using SDS Sample Buffer with 5% beta-Mercaptoethanol, followed by incubation for 10 min at 95°C. Using a magnetic separation rack the supernatant was collected and centrifuged at 10,000 x g at 4°C. The supernatant was loaded onto a 6% polyacrylamide gel and Western blotting was performed as outlined previously.
2. ***Mass spectrometry analysis***: 66% of sample was enzymatically digested and eluted as described previously [53]. In brief, samples was eluted using 60 µL of solution containing 2 M urea diluted with 50 mM Tris-HCL pH 7.5 containing 5 µg/mL Trypsin. Samples were incubated at 27°C for 30 min in a thermomixer at 800 rpm to aid digestion. A magnet was used to separate the beads from the supernatant. Beads were washed again with 25 µL of solution containing 2 M urea diluted in 50 mM Tris-HCL pH7.5 and 1mM DTT) and the supernatant was collected. This process was repeated twice. Peptides were incubated at RT overnight to facilitate digestion. 20 µL of iodoacetamide (5 mg/mL) was added to the samples and incubated for 30 min in the dark. To halt the reaction 1 μL of trifluoroacetic acid (100% TFA) (Sigma, Cat. no. 302031) was added to each sample and samples were desalted using C18 stagetips [54]. Peptides were eluted in 2.5% acetonitrile/0.5% acetic acid buffer. Peptides (400 ng -2,800 ng) were analysed using a timsTOF (Bruker) mass spectrometer. The mass spectrometry proteomics data have been deposited to the ProteomeXchange Consortium via the PRIDE [52] partner repository with the dataset identifier PXD080845 (to be made publicly accessible upon publication).

### Proteomic Analysis

Differentially expressed proteins (DEPs) were analysed using Ingenuity Pathway Analysis (Qiagen) and STRING.

### Statistical analysis and Data Visualisation

Data was visualised and analysed using GraphPad Prism (version 10.6.1 for Windows, GraphPad Software, Boston, Massachusetts USA). Data normality was assessed using a Shapiro-Wilk Test. For normally distributed data, statistical significance was determined using a one-way ANOVA followed by Sidak’s multiple comparisons test for pre-selected pairs. For non-parametric data, a Kruskal-Wallis test was applied with Dunn’s multiple comparisons test on pre-selected pairs. Data represent mean ± SEM unless otherwise stated. Proteomic data was analysed using a paired t-test with statistical (p=<0.05) and fold change (<-1.2 or >1.2) cut-offs applied. Metabolic flux analysis and angiogenesis assay time course was analysed using a linear mixed effects model including Group, Time, and Group × Time interaction, with well (technical replicates) included as a random effect and the estimated marginal means ± SE were graphed. Statistical comparisons between groups were performed at each time point using estimated marginal means with Holm correction for multiple comparisons. An asterisk (*) denotes p<0.05, while **=p<0.01, ***=p<0.001, ns = not statistically significant. Violin superplots were used to visualise metabolic flux analysis and imaging data. Large symbols represent the mean of each biological replicate, while smaller symbols represent individual technical replicates within each biological replicate.

## Results

Targeted knockdown of *NOTCH3* in HAoSMCs provides insights into the role of NOTCH3 in regulating VSMC function, phenotype and downstream signalling pathways. To investigate the role of NOTCH3 signalling in VSMCs, HAoSMCs were transfected with NOTCH3-targeting siRNA. Cells were examined under basal conditions and following stimulation with PDGF or TGF-β to model key signalling pathways that regulate VSMC biology. PDGF which has been demonstrated to increase NOTCH3 expression [55], was selected due to its established role in mural cell differentiation and recruitment [56, 57]. TGF-β was used as an inducer of smooth muscle actin, collagen and other ECM-associated genes [58, 59].

NOTCH3 siRNA reduced *NOTCH3* transcript expression under control, PDGF and TGF-β conditions by 87%, 83% and 85%, respectively, compared with negative control siRNA (Neg Ctrl siRNA) (**Fig. 1A**). Efficient suppression of NOTCH3 was confirmed at the protein level across all treatment conditions (**Fig. 1B-C**). Cell viability was unchanged between the NOTCH3 siRNA and the Neg Ctrl siRNA groups (**Fig. 1D**). Furthermore, although NOTCH3 has been implicated in regulating cell proliferation rates in cancer models [60], NOTCH3 suppression did not alter HAoSMC proliferation rates under the conditions examined (**Supplemental Fig. 4A-C**).

**Figure 1:**
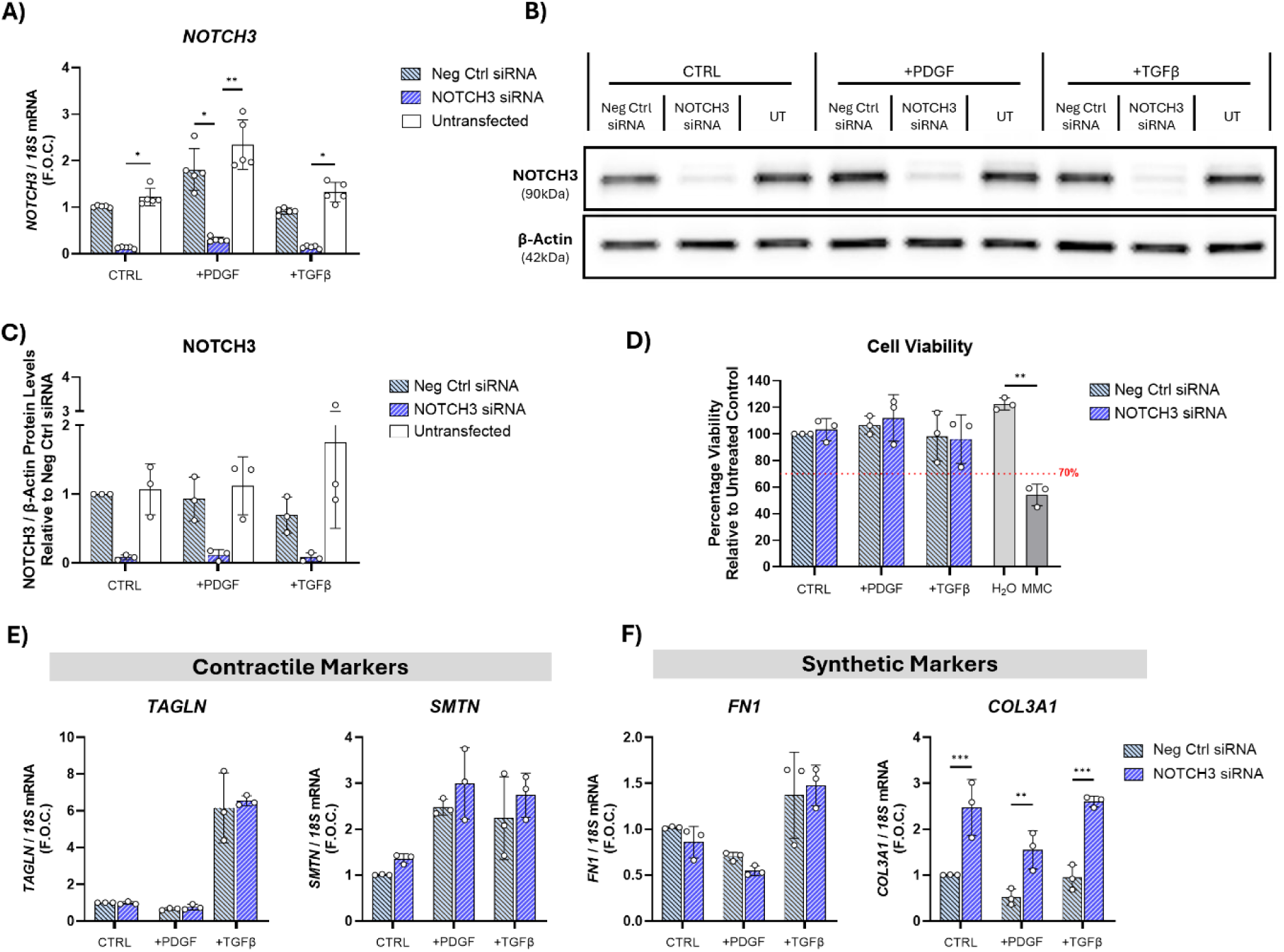
Knockdown of NOTCH3 in HAoSMCs increases the expression of *COL3A1*. HAoSMCs were treated with NOTCH3 siRNA or the negative control (Neg Ctrl) siRNA or left untransfected (UT) for 6 h followed by stimulation with PDGF (20ng/mL) and TGFβ (10ng/mL) or vehicle control (CTRL; PBS with 0.01% BSA) for 48 h. **A)** *NOTCH3* gene expression was analysed by qRT-PCR using 18S rRNA as the endogenous control. 5 independent experiments were performed (N=5). **B)** Representative Western blot of NOTCH3 protein, with β-actin as the control (N=3). **C)** Densitometry analysis of NOTCH3 protein levels normalized to β-actin and relative to Neg Ctrl siRNA (N=3). **E)** Cell viability was assessed after 48 h using an MTT assay and normalized to Neg CTRL siRNA. Mitomycin-C (MMC, 20 µg/mL) was included as a positive control (N=3). **E)** Gene expression of SMC contractile markers, *TAGLN and SMTN*, was assessed by qRT-PCR using 18S rRNA as the endogenous control. **F)** Gene expression of SMC synthetic markers, *FN1* and *COL3A1*, was assessed by qRT-PCR using 18S rRNA as the endogenous control (N=3). Statistical analysis was performed using (i) a Kruskal-Wallis test with Dunn’s multiple comparisons test (A-C) and (ii) an ordinary one-way ANOVA with Sidak’s multiple comparisons test (D-F). *=p<0.05, **=p<0.01, ***=p<0.001.

Next, to investigate the effects of NOTCH3 knockdown on the HAoSMC phenotype, established markers of SMC contractile, synthetic and inflammatory states were assessed. Expression of contractile markers, Transgelin (*TAGLN*) and Smoothelin (*SMTN*) remained unchanged following NOTCH3 suppression (**Fig. 1E**). The inflammatory markers, Interleukin-6 (*IL6*) and Nuclear Factor Kappa B Subunit 1 (*NFKB1*), were also unchanged following NOTCH3 knockdown (**Supplemental Fig.4D-E**). In contrast, the synthetic marker, Collagen Type III Alpha 1 Chain (*COL3A1*), was significantly increased across all conditions, whereas Fibronectin 1 *(FN1*) expression was unaffected (**Fig. 1F**). *COL3A1* encodes a major component of fibrillar collagen found in tissues such as blood vessels, bowel and skin, where it contributes to structural integrity, and plays a role in wound healing [61, 62]. Collectively, although siRNA-mediated suppression of NOTCH3 does not induce a broad transition towards a contractile, synthetic, or inflammatory phenotype, it selectively alters ECM-related gene expression through increased *COL3A1* expression.

### NOTCH3 maintains extracellular matrix modelling and metabolic pathways in HAoSMCs

To further assess the broader cellular impact of siRNA-mediated loss of NOTCH3 in HAoSMCs, mass spectrometry-based proteomic analysis was performed. Differentially expressed proteins (DEPs) were identified by comparing NOTCH3 siRNA-treated cells with Neg Ctrl siRNA-treated cells under basal (CTRL), PDGF and TGF-β conditions.

Under basal conditions, NOTCH3 knockdown increased the ECM-associated proteins Collagen Type V Alpha 2 Chain (COL5A2), Fibrillin 1 (FBN1) and Laminin Subunit Beta 2 (LAMB2) and also increased actin cytoskeleton-associated proteins, Formin Binding Protein 1 (FNBP1), Cortactin (CTTN) and LIM Domain Containing Preferred Translocation Partner In Lipoma (LPP) (**Fig. 2A**). Following PDGF-stimulation, NOTCH3 knockdown increase several ECM-associated proteins including multiple collagens (COL3A1, COL6A3, COL7A1, COL12A1), Secreted Protein Acidic And Cysteine Rich (SPARC), Serpin Family H Member 1 (SERPINH1) and latent TGF-β binding protein 1 (LTBP1), proteins typically associated with a synthetic SMC phenotype that is proliferative and migratory (**Fig. 2B**). Notably, LTBP1 and FBN1 have previously been identified within NOTCH3-ECD deposits in CADASIL post-mortem brain vessels [26] and LTBP1 can directly bind to mutant NOTCH3 [63]. Consistent with proteomics findings, LTBP1 gene expression showed a trend towards increased expression following NOTCH3 knockdown (**Supplemental Fig. 5A**). Following TGF-β stimulation, NOTCH3 knockdown increased actin cytoskeletal-associated proteins CTTN, Drebrin 1 (DBN1) and Diaphanous Related Formin 1 (DIAPH1), Myotrophin (MTPN) and Myosin Phosphatase Rho Interacting Protein (MRIP), alongside an increase in Matrix Metallopeptidase 14 (MMP14) (**Fig. 2C**). Collectively, multiple glycolysis-associated proteins where decreased while several of the most highly upregulated proteins were involved in regulating actin cytoskeletal remodelling, cell shape, focal adhesion dynamics and cell motility.

**Figure 2:**
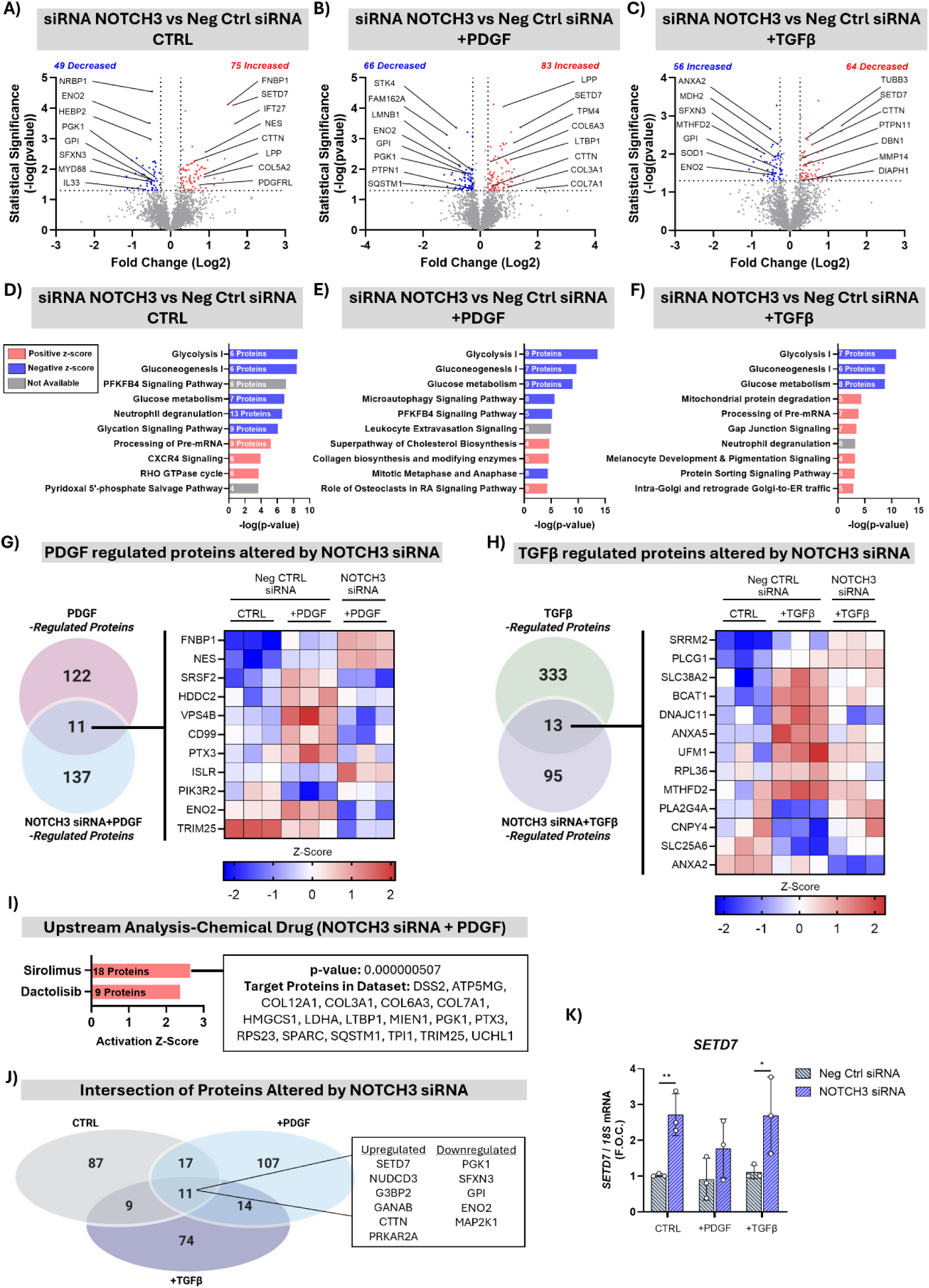
Mass spectrometry-based proteomic analysis of the effects of NOTCH3 knockdown in HAoSMCs. HAoSMCs were treated with NOTCH3 siRNA for 6 h followed by stimulation with PDGF (20ng/mL) and TGF-β (10ng/mL) or vehicle control (CTRL; PBS with 0.01% BSA) for 48 h. **A-C)** Scatter plots showing differential protein expression following NOTCH3 siRNA treatment in comparison to Neg Ctrl siRNA in the presence of **A)** CTRL, **B)** PDGF, and **C)** TGF-β. Thresholds were set at a fold change cut off of ± 1.2 and statistical significance at p<0.05. Several increased (red) and decreased (blue) proteins of interest were labelled. **D-F)** Canonical pathway analysis performed on each dataset using IPA, filtered by statistical significance (-log(p-value)). **G)** Venn diagram illustrating the intersection between PDGF regulated proteins and NOTCH3 siRNA + PDGF-regulated proteins, accompanied by a heatmap of z-scores for the 11 overlapping proteins. **H)** Venn diagram illustrating the intersection between TGF-β-regulated proteins and NOTCH3 siRNA + TGF-β-regulated proteins, accompanied by a heatmap of z-scores for the 13 overlapping proteins. **I)** IPA upstream regulator analysis of the differentially altered proteins from NOTCH3 siRNA + PDGF treatment, filtering for ‘Chemical Drugs’. **J)** Venn diagram illustrating the intersection of the proteins altered by NOTCH3 siRNA across all three conditions (CTRL, PDGF, TGF-β) highlighting the 11 commonly altered proteins. **K)** *SETD7* gene expression analysed by qRT-PCR using *18S rRNA* as the endogenous control (N=3). Statistical analysis was performed using a ordinary one-way ANOVA with Sidak’s multiple comparisons test. *=p<0.05. **=p<0.01.

Ingenuity Pathway Analysis (IPA) of proteins altered by NOTCH3 siRNA predicted a consistent inhibition of ‘Glycolysis I’ and ‘Glucose metabolism’ across all treatment conditions, reflected by the significant negative activation z-scores. Notably, the ‘Collagen biosynthesis and modifying enzymes’ pathway was predicted to be activated specifically in PDGF-stimulated cells following NOTCH3 knockdown (**Fig. 2D-F**).

As PDGF and TGF-β did not significantly alter NOTCH3 protein abundance, proteins regulated by each growth factor were first identified and subsequently compared with those altered following NOTCH3 knockdown under the corresponding stimulation conditions to determine how loss of NOTCH3 modifies growth factor responses. Eleven proteins overlapped between the PDGF and NOTCH3 siRNA + PDGF datasets. FNBP1, Nestin (NES) and Immunoglobulin Superfamily Containing Leucine Rich Repeat (ISLR), which were reduced by PDGF stimulation, were increased following NOTCH3 knockdown. Conversely, the PDGF-induced upregulation of CD99 and Pentrxin-3 (PTX3) was attenuated by NOTCH3 knockdown. Similarly the glycolytic enzyme, Enolase 2 (ENO2) was also decreased with NOTCH3 siRNA (**Fig. 2G**). TGF-β treatment altered 346 proteins, of which 13 were also regulated by NOTCH3 knockdown. The TGF-β-induced mitochondrial proteins, DNAJC11 and MTHFD2, were attenuated following NOTCH3 knockdown. In addition, the signalling proteins Annexin A2 (ANXA2), Phospholipase C Gamma 1 (PLCG1), and Phospholipase A2 Group IVA (PLA2G4A), which have established roles in angiogenesis [64, 65], were also dysregulated (**Fig. 2H**).

IPA upstream regulator analysis, filtered for ‘Chemical Drugs’, identified Sirolimus (Rapamycin) as the highest ranking predicted upstream regulator, targeting multiple collagen-associated proteins and LTBP1 (**Fig. 2I**). This finding is notable given that Sirolimus (Rapamycin) promotes the vascular SMC contractile phenotype while suppressing collagen synthesis via inhibition of mTOR signalling [66, 67].

Finally, proteins commonly dysregulated by NOTCH3 knockdown across all treatment conditions were identified. The glycolytic enzymes, ENO2 and Glucose-6-Phosphate Isomerase (GPI) were commonly decreased, whereas CTTN, a regulator of actin dynamics and cell migration, was consistently increased. Among the most highly upregulated proteins was SET Domain Containing 7, Histone Lysine Methyltransferase (SETD7) (**Fig. 2J; Supplemental Fig. 5B**). SETD7 is a regulator of TGF-β1 activation in renal fibroblasts, specifically regulating SMAD activity [68] and has been posited as a promising target for pharmacological inhibition to treat conditions such as diabetes [69]. As SETD7 potentiates TGF-β signalling contributing to ECM gene expression and fibrosis [70] its increased levels following NOTCH3 knockdown were validated at the transcript level (**Fig. 2K**).

### Glycolysis and the glycolytic enzyme, ENO2, are reduced in HAoSMCs following knockdown of NOTCH3

To further investigate the metabolic changes induced by NOTCH3 knockdown, an interaction network of glycolysis-associated proteins altered across all treatment conditions was generated from the proteomic dataset (**Fig. 3A**). Consistent with the inhibition of glycolysis predicted by IPA, the majority of glycolytic proteins were decreased.

**Figure 3:**
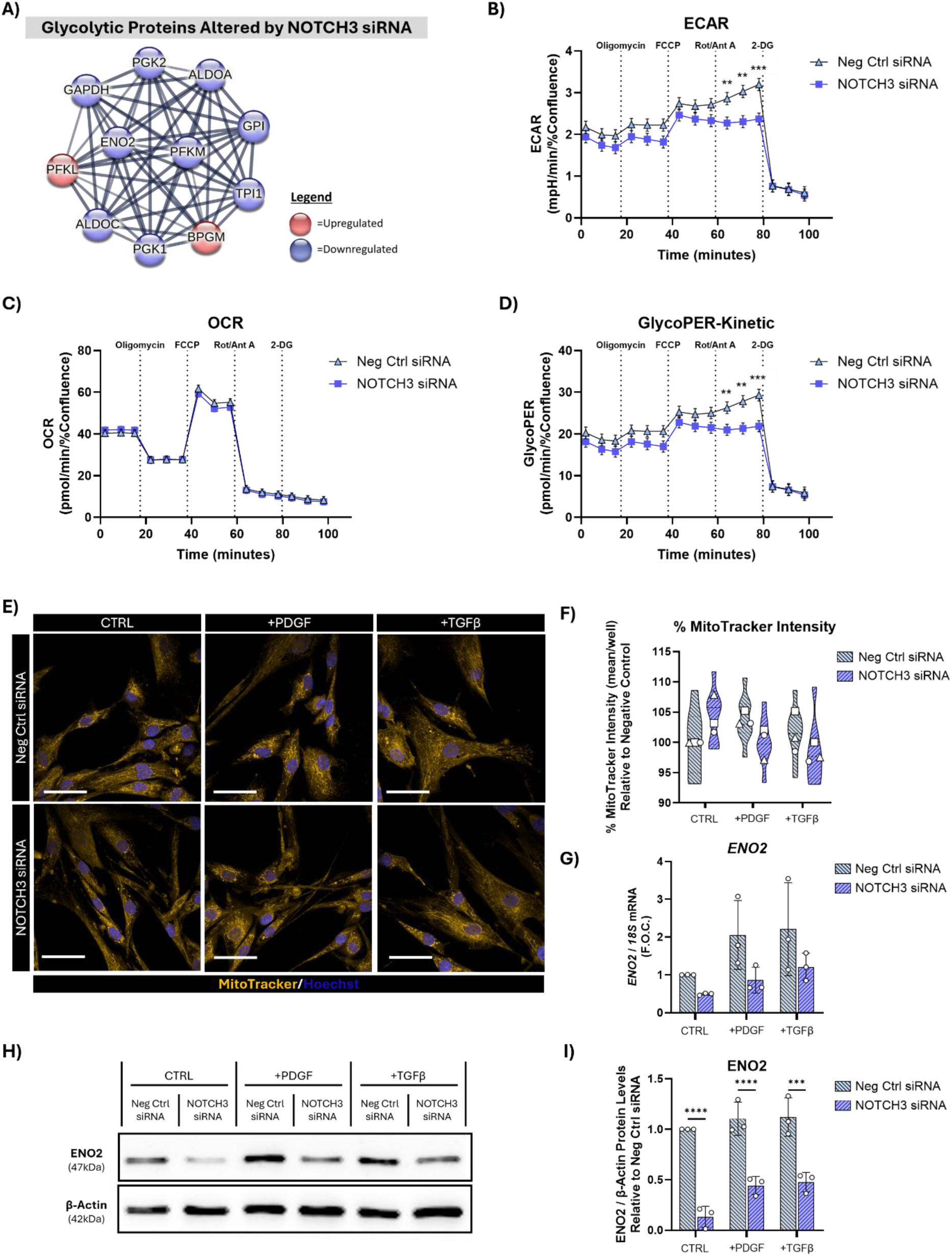
Glycolysis and the glycolytic enzyme, ENO2, are impaired in HAoSMCs following knockdown of NOTCH3. **A)** STRING interaction network of the glycolytic proteins altered by NOTCH3 siRNA identified by mass-spectrometry. **B-D)** HAoSMCs were treated with NOTCH3 siRNA for 6 h followed by a rest period of 24 h. Cells were then seeded into a Seahorse XF96 Cell Culture Microplate and allowed to adhere overnight. **A)** The extracellular acidification rate (ECAR). **C)** The oxygen consumption rate (OCR). **D)** Kinetic glycolytic proton efflux rate (glycoPER-kinetic). 3 independent experiments were performed with 8-9 technical replicates per condition (N=3, n=8-9). **E)** Representative images of mitochondria stained with MitoTracker and nuclei stained with Hoechst. **F)** Quantitative analysis of mitochondrial staining intensity using Harmony Software (N=3, n=3, 20 fields of view per well). **G)** *ENO2* gene expression analysed by qRT-PCR using *18S rRNA* as the endogenous control (N=3). **H)** Representative Western blot of ENO2 protein levels, with β-actin as the loading control (N=3). **I)** Densitometry analysis of ENO2 protein levels normalised to β-actin (N=3). For B-D a linear mixed-effects model was fitted to the data and values depict the estimated marginal means ± SE (N=3, n=8-9). Statistical comparisons between groups were performed at each time point using estimated marginal means with Holm correction for multiple comparisons. For F,G, and I, statistical analysis was performed using an ordinary one-way-ANOVA with Sidak’s multiple comparisons test. *=p<0.05. **=p<0.01, ***=p<0.001.

To functionally validate these findings, real-time metabolic flux analysis was performed in HAoSMCs following NOTCH3 knockdown. NOTCH3 siRNA decreased the extracellular acidification rate (ECAR), indicative of impaired glycolytic activity (**Fig. 3B**) while the oxygen consumption rate (OCR) was unchanged (**Fig. 3C**). This effect was most evident following inhibition of mitochondrial oxidative phosphorylation with Rotenone and Antimycin A, where negative control siRNA-treated cells exhibited the expected compensatory increase in glycolysis/ECAR that was attenuated following NOTCH3 knockdown. Consistent with the ECAR measurements, Glycolytic Proton Efflux Rate (glycoPER-kinetic), a more accurate and quantitative measurement of glycolytic flux, also showed reduced glycolytic capacity following NOTCH3 knockdown (**Fig. 3C**). A decreased trend was noted in ATP production, basal glycolysis and compensatory glycolysis with NOTCH3 knockdown (**Supplemental Fig. 6A-C**).

Previous studies reported structural and functional mitochondrial abnormalities in CADASIL-derived cells and tissues [71–73]. Therefore, mitochondrial morphology was assessed following NOTCH3 knockdown. Mitochondria number, morphology and fluorescence intensity were unchanged (**Fig. 3E-F**), suggesting that the metabolic deficits were not attributable to overt mitochondrial alterations.

Finally, ENO2, one of the glycolytic enzymes consistently reduced in the proteomic analysis, was validated at both the transcript and protein levels (**Fig. 3G-I**). ENO2 catalyses the conversion of 2-phosphoglycerate (2PG) into phosphoenolpyruvate (PEP) during glycolysis and its depletion has previously been shown to reduce ECAR [74]. Collectively, these results show that NOTCH3 knockdown impairs glycolytic metabolism in HAoSMCs without detectable changes in mitochondrial morphology, coinciding with reduced ENO2.

### NOTCH3 knockdown induces a VEGF-associated proteomic signature without altering endothelial angiogenesis

PANTHER pathway analysis of all differentially expressed proteins (DEPs) identified enrichment of proteins associated with angiogenesis and VEGF signalling (**Fig. 4A**), pathways previously reported to be dysregulated in CADASIL [43, 75]. Interactions among VEGF-associated proteins were visualised using STRING (**Fig. 4B**). Providing further mechanistic insight, IPA network analysis predicted inhibition of VEGF signalling following NOTCH3 knockdown (**Supplemental Fig. 7A**). Supporting this prediction, *VEGFA* expression was reduced following NOTCH3 knockdown in PDGF-and TGF-β-stimulated conditions (**Fig. 4C**) whereas Thrombospondin-1 (*THBS1*) expression was unchanged (**Supplemental Fig. 7B**).

**Figure 4:**
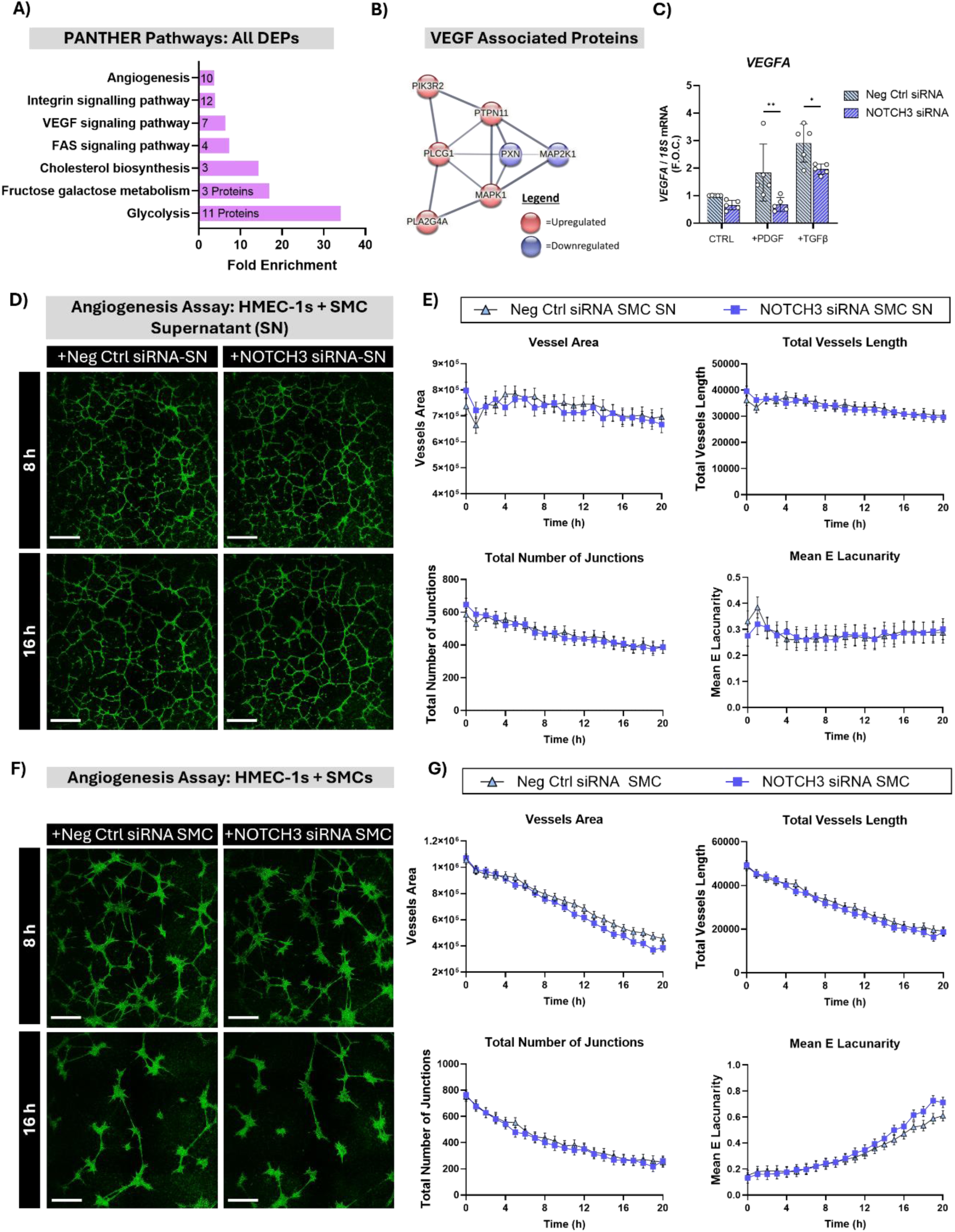
NOTCH3 knockdown alters angiogenic-associated proteins without affecting HAoSMC-supported endothelial network formation. **A)** PANTHER Pathway analysis of all differentially expressed proteins (DEPs) across NOTCH3 siRNA proteomic datasets, filtered for fold enrichment. **B)** STRING interaction network of VEGF signalling pathway-associated proteins. C) *VEGFA* gene expression analysed by qRT-PCR using *18S rRNA* as the endogenous control in HAoSMCs treated with NOTCH3 siRNA or negative control (Neg Ctrl) siRNA for 6 h, rested overnight, and stimulated with PDGF (20 ng/mL) or TGF-β (10 ng/mL) for 48 h (N=5). **D)** Representative images of HMEC-1 tube formation on Geltrex using 100 µL of conditioned supernatant from NOTCH3 siRNA-and Neg Ctrl siRNA-treated HAoSMCs captured by the Incucyte system after 8 and 16 h. Images were obtained over a time course of 20 h and pseudocoloured green for visualisation purposes and analysed using AngioTool2.0. **E)** AngioTool2.0 quantitative analysis of network parameters from condition supernatant experiments over a time course of 20 h (N=4, n=3). **F)** Representative images of co-cultured HAoSMCs (treated with NOTCH3 siRNA or Neg Ctrl for 6h and rested overnight) and HMEC-1 on Geltrex, captured by the Incucyte system after 8 and 16 h. Images were obtained over a time course of 20 h and pseudocoloured green for visualisation purposes and analysed using AngioTool2.0. **G)** AngioTool2.0 quantitative analysis of network parameters from co-culture assays over a period of 20 h (N=4, n=3). Scale bar = 800µm. For C, statistical analysis was performed using an ordinary one-way ANOVA with Sidak’s multiple comparisons test (N=3). For E and G, a linear mixed effects model was fitted to the data and values depict the estimated marginal means ± SE (N=4, n=3). Statistical comparisons between groups were performed at each time point using estimated marginal means with Holm correction for multiple comparisons. *=p<0.05, **=p<0.01.

To determine whether these molecular changes influenced endothelial angiogenesis, HMEC-1 cells were treated with conditioned media from unstimulated HAoSMCs following NOTCH3 knockdown or directly cultured with NOTCH3-deficient HAoSMCs. Although NOTCH3 knockdown was associated with a proteomic signature consistent with dysregulated angiogenic signalling, conditioned media (supernatant) from NOTCH3-knockdown HAoSMCs did not affect endothelial network formation in HMEC-1 cells (**Fig. 4D-E**).

As mural cells also support endothelial cells through direct cell-cell interactions HAoSMCs treated with NOTCH3 siRNA were co-cultured with HMEC-1 cells to assess endothelial network formation. NOTCH3 knockdown did not alter endothelial network formation in the co-culture model (**Fig. 4F-G**). Furthermore, NOTCH3 knockdown did not affect the ability of HAoSMC to form interconnected cellular networks in the absence of endothelial cells (**Supplemental Fig. 8A-B).** Collectively, these findings demonstrate that although NOTCH3 knockdown reduced VEGFA expression and was associated with a proteomic signature indicative of altered angiogenic signalling, these molecular changes were not accompanied by detectable alterations in endothelial network formation under the conditions examined.

### NOTCH3 knockdown remodels the actin cytoskeleton without altering HAoSMC migration

To further investigate the effects of NOTCH3 knockdown on HAoSMCs, DEPs were analysed by the Human Gene Atlas via ENRICHR which identified ‘Smooth Muscle’ as the top enriched term (**Supplemental Fig. 9A-C**). Interactions among ‘Smooth Muscle’-associated proteins altered by NOTCH3 siRNA were visualised using STRING (**Fig. 5A**). Cortactin (CTTN), a regulator of actin cytoskeleton dynamics, was identified as both a SMC-associated protein and a central hub within the IPA interaction network analysis (**Supplemental Fig. 9D**). IPA was used to identify DEPs involved in ‘Actin Cytoskeleton Signalling’ which included DIAPH1, and their interaction visualized using STRING (**Fig. 5B**).

**Figure 5:**
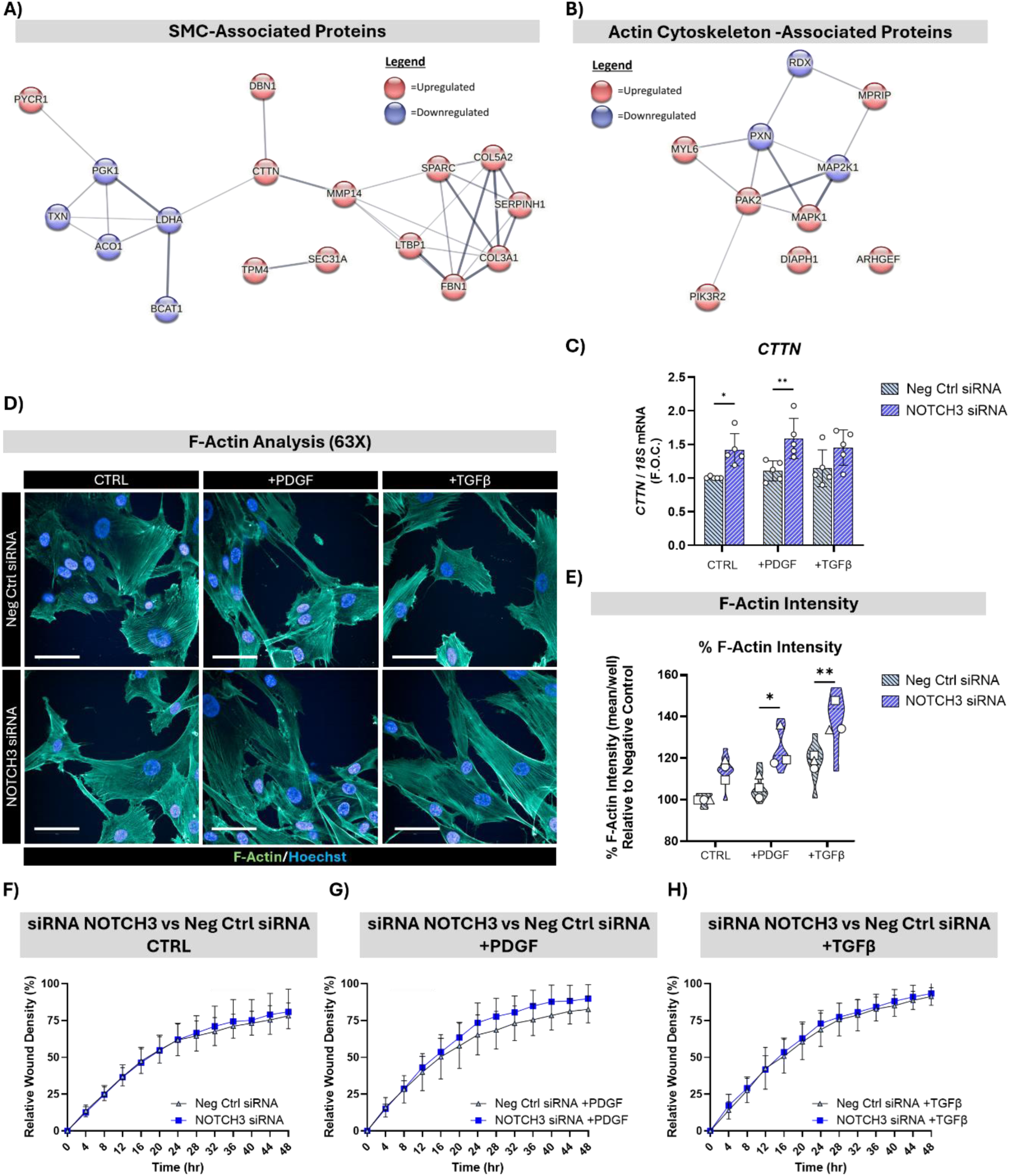
SMC-and actin-associated proteins are increased by knockdown of NOTCH3 in HAoSMCs. HAoSMCs were treated with NOTCH3 siRNA for 6 h, followed by stimulation with PDGF (20 ng/mL) and TGF-β (10 ng/mL) or vehicle control (CTRL; PBS with 0.01% BSA) for 48 h. **A)** STRING interaction network of SMC-associated proteins identified via Human Gene Atlas analysis (accessed using ENRICHR Software) of the NOTCH3 siRNA proteomic dataset. **B)** STRING network visualisation of actin cytoskeleton signalling-associated proteins identified from IPA analysis of the NOTCH3 siRNA proteomic dataset. **C)** *CTTN* gene expression was analsyed by qRT-PCR using *18S rRNA* as the endogenous control (N=5). **D)** Representative 63X confocal microscopy images of treated HAoSMCs stained for F-Actin and Hoechst, captured on an Opera Phenix microscope and analysed using Harmony Software. **E)** F-Actin fluorescence intensity normalised to the Neg Ctrl siRNA with PBS (N=3, n=3, 20 fields of view per well). The violin plots depict the distribution of the data across technical replicates. **F-H)** Scratch wound assay assessing the impact of NOTCH3 siRNA on cell migration/wound closure under **F)** vehicle (CTRL), **G)** PDGF and **H)** TGF-β conditions. Cells were treated with NOTCH3 siRNA for 6 h, rested overnight, and pre-incubated with Mitomycin-C (20 µg/mL) for 2 h. A scratch was generated using the Incucyte WoundMaker and cells were stimulated with PDGF (20 ng/mL) and TGF-β (10 ng/mL) or vehicle control (CTRL; PBS with 0.01% BSA) for 48 h. Cells were analysed every 4 h for a total of 48 h (N=3, n=3). Scale bar = 50 µm. Statistical analysis of the means was performed using an ordinary one-way ANOVA with Sidak’s multiple comparisons test. *=p<0.05, **=p<0.01.

Consistent with the proteomic analysis, *CTTN* was increased at the transcript level following NOTCH3 knockdown (**Fig. 5C**). As CTTN directly binds and stabilises F-actin, promoting cytoskeletal remodelling processes involved in cell migration and invasion [76], and DIAPH1 is required for the assembly of F-Actin structures, the increase in these actin-associated proteins following NOTCH3 knockdown suggested altered actin dynamics. Confocal microscopy analysis confirmed increased F-Actin intensity following NOTCH3 knockdown (**Fig. 5D-E**).

To determine whether these cytoskeletal changes altered cell motility, a wound-healing assay was performed to assess cell migration. NOTCH3 knockdown did not affect relative wound density across the conditions (**Fig. 5F-H**). In parallel, the NOTCH inhibitor DAPT did not affect this process, however, as expected, Mitomycin C and Rotenone effectively suppressed relative wound density and cell proliferation (**Supplemental Fig. 10A-F**). Collectively, these results demonstrate that NOTCH3 knockdown remodels the actin cytoskeleton and alters SMC-associated proteins without producing detectable changes in HAoSMC migration under the conditions examined.

### NOTCH3 suppresses extracellular matrix remodelling in HAoSMCs

The connection between NOTCH3 and ECM regulation is well documented [25, 35, 77]. Notably multiple collagen subtypes including COL1A1, COL1A2, COL6A1, COL6A2, COL6A3 and COL12A1 accumulate within affected blood vessels in CADASIL [26]. IPA of the DEPs following NOTCH3 knockdown in PDGF-stimulated cells identified a prominent collagen interaction network containing the increased collagens alongside predicted activation of PDGF, AKT and collagen type IV (**Fig. 6A**). Given that *COL3A1* expression was identified as consistently increased following NOTCH3 knockdown across all 3 conditions, STRING analysis was performed to identify interacting DEPs, revealing ISLR and FBN1 as COL3A1-associated proteins (**Fig. 6B**).

**Figure 6:**
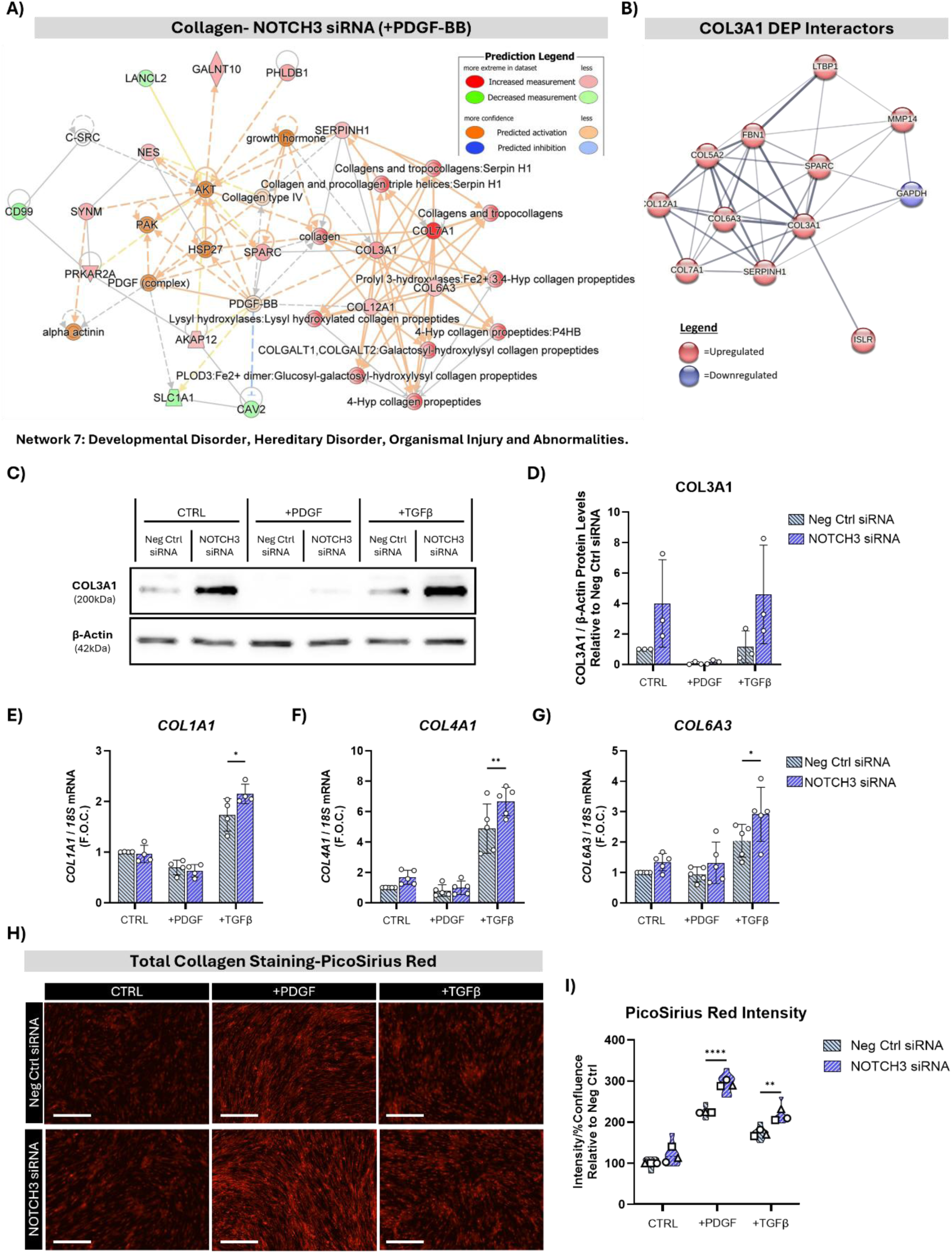
Collagen is increased following knockdown of NOTCH3 in HAoSMCs. HAoSMCs were treated with NOTCH3 siRNA or negative control (Neg Ctrl) siRNA for 6 h, followed by stimulation with PDGF (20 ng/mL) and TGF-β (10 ng/mL) or vehicle control (CTRL; PBS with 0.05% BSA) for 48 h prior to proteomic analysis was performed. **A)** IPA network analysis of DEPs altered by NOTCH3 siRNA in the presence of PDGF. Collagen was predicted to be increased in Network 7 titled ‘Developmental Disorder, Hereditary Disorder, Neurological Disease, Organismal Injury and Abnormalities’. **B)** STRING interaction network illustrating DEPs with known interactions with COL3A1. **C)** Representative Western blot of COL3A1 protein, with β-actin as the loading control (N=3). **D)** Densitometry analysis of Western blots for COL3A1, normalized to β-actin and relative to Neg Ctrl siRNA-CTRL (N=3). **E-G)** Gene expression of **E)** *COL1A1*, **F)** *COL4A1*, **G)** *COL6A3* analysed by qRT-PCR using *18S rRNA* as the endogenous control (N=4-5). **H)** Representative images of collagen stained with PicoSirius Red and captured on the Incucyte system after 4 days of stimulation with PDGF-BB, TGF-β or vehicle control (CTRL) in NOTCH3 siRNA-and Neg Ctrl siRNA-treated HAoSMCs. Scale bar = 400 µm. **I)** Quantitative analysis of PicoSirius Red fluorescence intensity normalized to cell confluence (N=3, n=3, 4 fields of view per well). The violin plots depict the distribution of the data across technical replicates. Statistical analysis was performed on the means using an ordinary one-way ANOVA with Sidak’s multiple comparisons test. *=p<0.05, **=p<0.01, ****<p<0.0001.

Consistent with gene expression analysis, COL3A1 protein levels were increased following NOTCH3 knockdown, with the greatest abundance observed under basal and TGF-β-stimulated conditions (**Fig. 6C-D**). In addition, TGF-β stimulation increased expression of other collagens and NOTCH3 knockdown further increased *COL1A1*, *COL4A1* and *COLcA3* expression in TGF-β-stimulated cells (**Fig. 6E-G**).

To independently validate these molecular changes, total collagen deposition was quantified by Picosirius Red staining following four days of PDGF-or TGF-β-stimulation post NOTCH3 knockdown (**Supplemental Fig. 11A**). Both PDGF and TGF-β stimulation increased collagen deposition, which was further enhanced by NOTCH knockdown (**Fig. 6H-I**). Although total collagen content increased, mature collagen fibrils were not detected under polarised light (data not shown).

Collectively these findings demonstrate that NOTCH3 knockdown promotes collagen expression and ECM deposition in HAoSMCs, which was most evident under growth factor stimulation.

### The NOTCH3 interactome in HAoSMCs comprises SMC-associated proteins, cytoskeletal regulators and ECM proteins

To define the endogenous NOTCH3 interactome in healthy HAoSMCs, NOTCH3 was immunoprecipitated using an antibody-based pull-down approach followed by mass spectrometry analysis. Immunoprecipitation of NOTCH3 was confirmed by Western blotting, with NOTCH3 detected at its expected molecular weight (∼90 kDa) in both whole-cell lysate and the NOTCH3 immunoprecipitate (**Fig. 7A**). Although non-specific binding was observed in the IgG control condition, mass spectrometry analysis confirmed the complete absence of NOTCH3 in the IgG control sample whereas NOTCH3 was markedly enriched in the target immunoprecipitation fraction (**Fig. 7B**).

**Figure 7:**
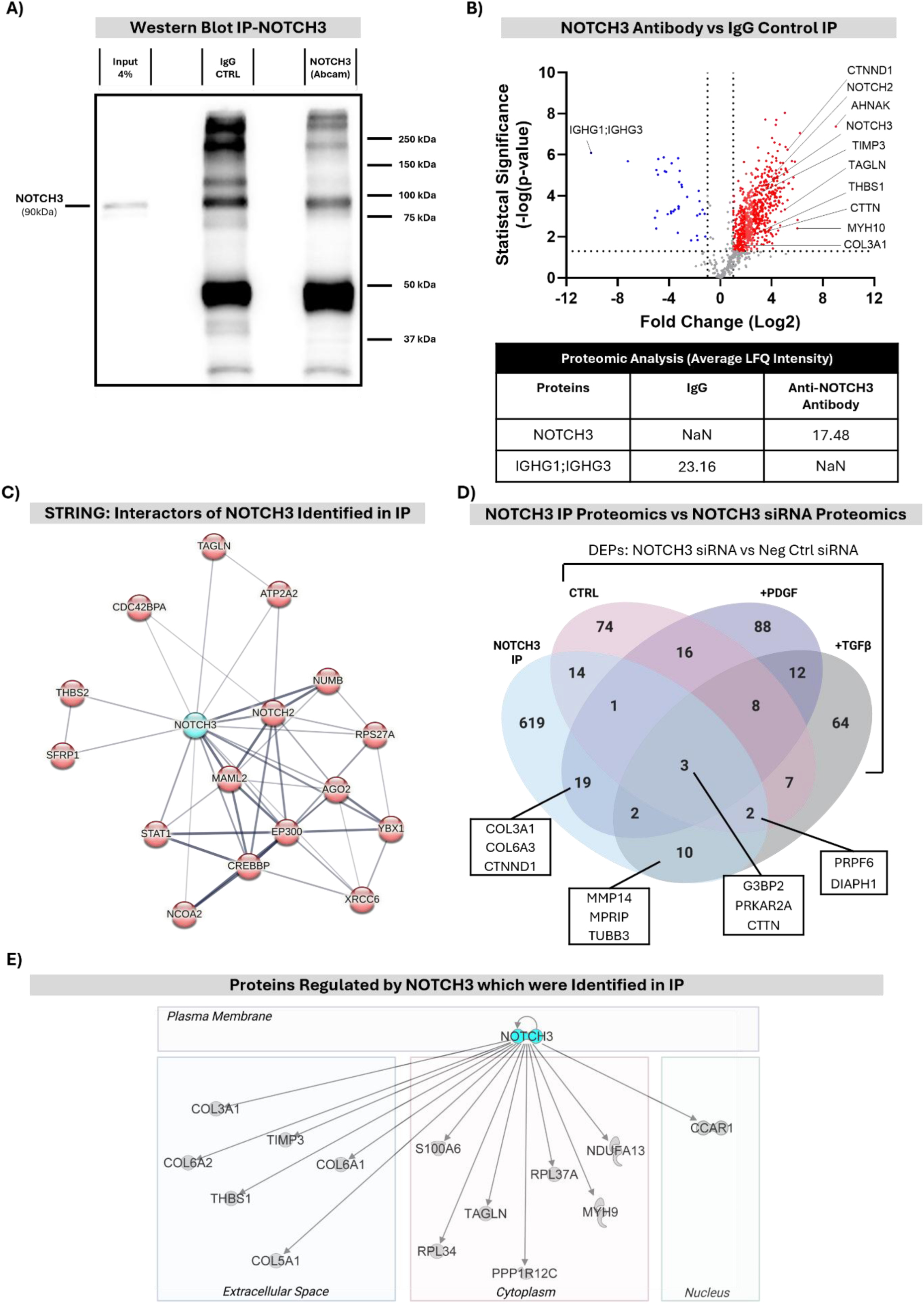
Identification of the protein interactome of NOTCH3 in HAoSMCs. **A)** HAoSMCs were seeded in a 10-cm plate and cultured for 48 h. Endogenous NOTCH3 was immunoprecipitated (IP) using an anti-NOTCH3 antibody (Abcam), with rabbit IgG used as the negative control. Immunoprecipitates were analysed by Western blot alongside input lysates (4% of the original total protein input) (N=4). **B)** Mass spectrometry-based proteomic analysis of NOTCH3 and IgG immunoprecipitates (N=4). The fold change of NOTCH3 immunoprecipitates was calculated relative to the IgG Control. Thresholds for interactors were defined as fold change (FC) >1.2 and statistical significance at p<0.05. Average label-free quantification (LFQ) intensity values for NOTCH3 and immunoglobulin heavy constant gamma 1/3 (IGHG1; IGHG3) are tabulated (NaN = not a number/below detection limit). **C)** STRING interaction network of known NOTCH3 interactors that were co-immunoprecipitated with NOTCH3. **D)** Venn diagram showing the intersection between the NOTCH3 protein interactome (IP-MS) and the DEPs from the NOTCH3 siRNA proteomic dataset. **E)** Overlay of molecules predicted to be regulated by NOTCH3 (via IPA upstream regulator analysis) with direct physical interactors identified by NOTCH3 IP-MS.

Proteomic analysis identified several NOTCH3-interacting proteins associated with collagen and ECM organisation which included COL3A1, COL5A1, COL6A1, COL6A2, COL6A3, Lysly Oxidase (LOX), Peroxidasin (PXDN), THBS1, THBS2, TIMP3 and MMP14. In addition, several cytoskeletal proteins were identified which included CTTN, Calponin-2 (CNN2), Transgelin (TAGLN), DIAPH1, Filamin-A (FLNA) as well as multiple myosin and tubulin family members. STRING analysis was performed to visualise known interactions between NOTCH3 and proteins identified in the immunoprecipitation dataset (**Fig. 7C**). Notably, the analysis identified both nuclear co-activator Mastermind-like Protein 2 (MAML2) and NUMB, established binding partners of the NOTCH3 ICD [78, 79], providing support for the specificity of the interactome. Integration of the NOTCH3 interactome with DEPs following NOTCH3 knockdown identified CTTN as both a NOTCH3-interacting protein and a downstream target of NOTCH3 signalling (**Fig. 7D**). Additional proteins identified at the intersection of these datasets were COL3A1, MMP14, MPRIP and DIAPH1. Furthermore intersection of this NOTCH3 IP data with proteomic analysis of CADASIL blood vessels [26] identified several overlapping ECM-associated proteins including TIMP3, COL6A1, COL6A2 and COL6A3 (**Supplemental Fig. 12A**).

IPA upstream regulator analysis of the NOTCH3 immunoprecipitation dataset identified TGF-β1 as a significant predicted upstream regulator, supporting convergence between NOTCH3 and TGF-β signalling pathways (**Supplemental Fig. 12B**). Among ion channels, Potassium Inwardly Rectifying Channel Subfamily J Member 2 (KCNJ2) emerged as a predicted upstream regulator (**Supplemental Fig. 12B**) and has previously been implicated in CADASIL murine models [80] and in patients who have deficits in functional hyperemia [81]. Finally, IPA upstream regulator analysis was filtered for NOTCH3 which generated a list of target molecules known to be regulated by NOTCH3, which were subsequently grouped according to their subcellular localisation. Several collagens including COL3A1 were identified as both proteins regulated by NOTCH3 signalling and components of the NOTCH3 interactome (**Fig. 7E**). Collectively, these findings demonstrate that NOTCH3 interacts with proteins involved in VSMC homeostasis, ECM regulation and cytoskeletal organisation, with several interactors also exhibiting altered expression following loss of NOTCH3.

## Discussion

In this study, we investigated the molecular and functional consequences of siRNA-mediated NOTCH3 knockdown in HAoSMCs to better define the physiological role of NOTCH3. NOTCH3 knockdown induced coordinated remodelling of ECM, cytoskeletal and metabolic pathways, characterised by increased collagen, Cortactin (CTTN) and F-actin abundance, alongside a reduction in glycolysis and ENO2. Despite these widespread molecular changes, key functional properties of VSMCs, including proliferation, migration and support of endothelial network formation, were largely preserved. Together, these findings identify NOTCH3 as a key regulator of VSMC homeostasis and highlights the complexity of targeting NOTCH3 signalling where modulation of specific pathways may occur without immediate loss of vascular cell function.

VSMCs are the principal source of ECM proteins within the tunica media, where collagens provide tensile strength and structural support to the vessel wall. Our findings suggest that NOTCH3 contributes to the maintenance VSMC ECM homeostasis by regulating collagen expression and modulating cellular responses to PDGF and TGF-β. The increased collagen production following NOTCH3 knockdown is particularly relevant to CADASIL, in which excessive collagen deposition and altered distribution are prominent pathophysiological features [25, 82, 83]. Increased accumulation of collagen types I, III, V and VI has been reported in CADASIL cerebral blood vessels [25]. Here, loss of NOTCH3 consistently increased *COL3A1* expression and total collagen. COL3A1 is a major fibrillar collagen that provides tensile strength and structural integrity to the vascular wall, and increased type III collagen deposition is a feature of several fibrotic diseases [61]. Notably, heterozygous mutations in *COL3A1* cause the vascular form of Ehlers-Danlos syndrome (vEDS), which is characterized by fragile blood vessels, with risk of aneurysm formation and arterial rupture, resulting from reduced or structurally abnormal type III collagen [84]. Although CADASIL and vEDS arise through distinct pathogenic mechanisms, both conditions highlight the importance of maintaining appropriate type III collagen homeostasis for vascular integrity. In addition to COL3A1, increased expression of *COL1A1*, *COL4A1*, and *COL6A3* following NOTCH3 knockdown was only observed in the presence of TGF-β. *COL4A1* exhibited the greatest induction following TGF-β stimulation that was further augmented by NOTCH3 knockdown. In contrast to our findings, NOTCH3 knockdown decreased *COL4A1* and overexpression of NOTCH3 ICD increased *COL4A1* in unstimulated lung fibroblasts [85]. Furthermore, *COL4A1* and other mural cell-associated genes were increased by overexpression of NOTCH3 ICD in neural crest cells [44]. These differences suggest that regulation of COL4A1 by NOTCH3 is context dependent and may vary according to cell type, activation state and surrounding cytokine milieu. Collectively, these finding suggests that NOTCH3 not only regulates basal COL3A1 homeostasis but also modulates the collagen response of VSMCs to profibrotic stimuli such as TGF-β.

Under physiological conditions, adult blood vessels have VSMCs that exist predominantly in a contractile state, characterised by low rates of proliferation, migration and ECM synthesis [3]. Although NOTCH3 knockdown altered collagen-and actin-associated proteins, it did not affect any of the classical markers of the contractile phenotype, including *TAGLN* and *SMTN*, nor did it alter VSMC proliferation or migration. While increased collagen synthesis is typically associated with a synthetic VSMC state, the absence of corresponding changes in contractile markers or cellular behaviour suggests that loss of NOTCH3 alone is insufficient to drive phenotypic switching. Instead, our findings indicate that NOTCH3 regulates specific aspects of ECM remodelling without inducing a global transition between contractile and synthetic VSMC states.

These observations are consistent with reports that CRISPR-Cas9-mediated NOTCH3 gene knockout in pericytes did not alter the expression of pericyte markers NG2, RGS5, PDGFRB, and CDH2 [86], suggesting that NOTCH3 is not essential for maintenance of mural cell identity. Furthermore, although TAGLN and SMTN expression are increased in cerebral vessels from patients with CADASIL [24], neither protein was altered following acute NOTCH3 knockdown, highlighting important differences between pathogenic NOTCH3 mutations and transient loss of NOTCH3 expression.

Despite preservation of the contractile phenotype, NOTCH3 knockdown promoted cytoskeletal remodelling, characterized by increased abundance of actin-associated proteins DBN1, DIAPH1, MTPN and MRIP, coupled with increased F-actin intensity and CTTN upregulation. CTTN, an F-actin binding protein, is a key regulator of actin polymerisation and cytoskeletal dynamics [87] that was also identified here as a NOTCH3 binding protein. CTTN represents a potential molecular link between NOTCH3 signalling and the regulation of actin dynamics that has been previously described [27, 88]. Consistent with our findings shRNA-mediated knockdown of NOTCH3 altered actin organisation in healthy VSMCs [27], while VSMCs derived from iPSCs of patients with CADASIL exhibited abnormal F-actin organisation and impaired contractility [89]. Notably, the cytoskeletal changes induced by NOTCH3 knockdown in the present study did not translate into altered cell migration. Together these findings suggest that NOTCH3 plays a more prominent role in regulating cytoskeletal architecture than in controlling VSMCs phenotypic switching.

Here we identified a previously unrecognised association between NOTCH3 and the histone lysine methyltransferase, SETD7, across all experimental conditions. The inverse relationship between NOTCH3 and SETD7 suggests a potential regulatory axis, although whether SETD7 functions as a downstream effector of NOTCH3 signalling or is induced as a compensatory response to NOTCH3 loss remains to be determined. SETD7 has been implicated in the regulation of TGF-β signalling, fibrosis and collagenase. Specifically, SETD7 potentiates TGF-β signalling by targeting the inhibitory SMAD7, thereby promoting the expression of ECM genes [70]. Conversely, inhibition of SETD7 increases SMAD7 and attenuates TGF-β signalling [70].

Given the increase in collagen expression following NOTCH3 knockdown, the accompanying upregulation of SETD7 may represent a compensatory mechanism to limit excessive ECM accumulation as SETD7 has also been reported to regulate collagenase gene transcription [90]. Although this hypothesis requires experimental validation, it suggests that SETD7 may contribute to maintaining vascular ECM homeostasis following disruption of NOTCH signalling. Beyond ECM regulation, SETD7 has been implicated in vascular remodelling as increased SETD7 promotes transcription of the anti-angiogenic factor Semaphorin-3G, whereas pharmacological inhibition of SETD7 enhances neovascularisation and perfusion in a murine model of diabetic hindlimb ischaemia [69]. Further work is required to elucidated the NOTCH3-SETD7 relationship in ECM remodelling and vascular adaptation.

Knockdown of NOTCH3 reduced *VEGFA* expression in PDGF-and TGF-β-stimulated HAoSMCs. VEGFA is a key regulator of endothelial cell survival, proliferation, migration, permeability and angiogenesis through activation of VEGFR1 and VEGFR2 signalling [91]. NOTCH3 has been previously linked to VEGFA and angiogenesis [92, 93]. Consistent with our findings, VEGFA expression and VEGFR2 levels were reduced in the microvasculature of Notch3-knockout mice following angiotensin II exposure [94], supporting a role for NOTCH3 in regulating VEGF signalling.

Previous studies have also demonstrated that iPSC-derived mural cells (iMCs) from patients with CADASIL exhibit reduced VEGF secretion and expression and fail to adequately support endothelial network formation in co-culture models [43, 89]. Notably, siRNA-mediated knockdown of NOTCH3 restored VEGF secretion and rescued EC network formation specifically in CADASIL iMCs, whereas no effect was observed in control iMCs [43]. Similarly, in our study there was no effect on angiogenic network formation following acute knockdown of NOTCH3 in healthy HAoSMCs. This is consistent with reports that CRISPR-Cas9-mediated deletion of NOTCH3 in pericytes does not alter vascular network formation *in vitro* [86]. The contrasting responses between healthy cells and CADASIL-derived mural cells further support the concept that pathogenic NOTCH3 mutations do not simply represent a loss of NOTCH3 function but instead confer altered or neomorphic signalling properties that disrupt vascular homeostasis.

NOTCH3 signalling has previously been implicated in the regulation of cellular metabolism and mitochondrial homeostasis. In VSMCs from patients with CADASIL, mutant NOTCH3 has been associated with increased mitochondrial number, accumulation of morphologically abnormal mitochondria, and a reduction in functional mitochondria [71]. Similarly, muscle biopsy specimens from patients with CADASIL contain enlarged mitochondria [73] while patient-derived myofibroblasts and fibroblasts exhibited reduced mitochondrial content and a fragmented mitochondrial network [72]. Here we identified no alterations in the mitochondrial number and size within these cells which suggest that loss of NOTCH3 does not impact the mitochondrial profile. In contrast, acute knockdown of NOTCH3 in HAoSMCs produced modest metabolic alterations. Proteomic pathway analysis predicted inhibition of glycolysis, which was supported by a reduction in the glycolytic enzyme, ENO2, and a modest decrease in ECAR. NOTCH3 knockdown impaired the ability of HAoSMCs to increase glycolytic flux in response to mitochondrial inhibition, indicating reduced metabolic flexibility. Despite these changes, basal and compensatory glycolysis were only modestly affected, and neither migration nor proliferation was impaired. Consistent with a broader role for NOTCH3 in cellular metabolism, shRNA-mediated knockdown of NOTCH3 decreased OXPHOS and glycolytic potential in cancer cells stimulated with Fulvestrant [95]. Together, these findings suggest that NOTCH3 contributes to the regulation of glycolytic capacity in VSMCs but is not essential for maintaining the metabolic activity required to support fundamental cellular functions under basal conditions.

To fully understand the physiological role of NOTCH3 in VSMCs, it is important to define the protein interactome of this transmembrane receptor. Immunoprecipitation of NOTCH3 followed by proteomic analysis identified numerous ECM, collagen and cytoskeletal-associated proteins. The identification of BAG2 and TUBB3 as NOTCH3-interacting proteins is consistent with previous studies [96, 97]. Together with the detection of established NOTCH3 interactors, MAML2 and NUMB [78, 79], these findings provide independent validation of our NOTCH3 interactome dataset. Intersection with the NOTCH3 knockdown proteomic data identified several proteins that were both interactors of NOTCH3 and increased following loss of NOTCH3. Notably, COL3A1, which was consistently upregulated following NOTCH3 knockdown, was also identified as a NOTCH3-interacting protein suggesting that NOTCH3 may influence collagen homeostasis through both physical association and downstream regulation.

Defining the physiological NOTCH3 interactome is particularly relevant in the context of CADASIL, where the mutant NOTCH3 extracellular domain aberrantly accumulates and binds extracellular matrix proteins within the vessel wall [35, 63, 98]. Our findings demonstrate that interactions between NOTCH3 and collagen proteins occur under normal physiological conditions in healthy VSMCs, suggesting that these interactions are not inherently pathological. Instead, CADASIL-associated NOTCH3 mutations may strengthen, stabilise or prolong these physiological interactions through altered binding properties of the mutant ECD. Combined with excessive collagen production and impaired clearance of mutant NOTCH3 ECD, this aberrant retention of physiological binding partners may contribute to the progressive vascular fibrosis and vessel wall pathology characteristic of CADASIL.

In this study, NOTCH3 knockdown did not alter VSMC proliferation, migration or endothelial network formation, suggesting that several similar functional abnormalities in CADASIL models [43, 89] may primarily result from mutation specific mechanisms rather than reduced NOTCH3 expression alone. CADASIL-associated NOTCH3 mutations may exert toxic gain-of-function effects through ECD aggregation [99], altered ligand interactions [100], and/or impaired clearance of the mutant receptor [101]. Furthermore, *NOTCH3* expression was reduced by approximately 85% rather than completely ablated, and the preservation of these core cellular functions may reflect residual NOTCH3 activity and/or compensatory signalling through redundant pathways.

In conclusion, siRNA-mediated knockdown of NOTCH3 recapitulated some features of CADASIL pathology, including increased ECM production, particularly collagen, and cytoskeletal remodelling. These findings suggest that disruption of normal NOTCH3 function contributes to aspects of vascular remodelling, whereas many of the functional abnormalities observed in CADASIL are likely driven by mutation-specific gain-of-function mechanisms associated with the mutant NOTCH3 ECD.

Our findings highlight important considerations for therapeutic strategies aimed at modulating NOTCH3 signalling. Although siRNA-mediated reduction of NOTCH3 could theoretically reduce ECD accumulation and subsequent GOM formation that contribute to CADASIL pathology, our data suggest that sustained NOTCH3 suppression may also induce broader changes in VSMC biology, including alterations in collagen expression, VEGFA regulation, and glycolytic capacity. Given that excessive ECM deposition is a prominent downstream feature of CADASIL pathology [26, 83], therapeutic approaches resulting in prolonged NOTCH3 suppression may require careful consideration regarding the timing, duration, and extent of pathway inhibition. Given our findings in cells and prior reports of abnormal molecular phenotypes in Notch3 knockout mouse models, diseases caused by NOTCH3 mutations may not reflect a simple toxic gain-of-function mechanism, and allele-specific knockdown of mutant transcripts warrants consideration.

A knockdown-replacement strategy, combining transient NOTCH3 suppression with restoration of functional NOTCH3 expression through mRNA replacement, may also represent a potential approach to reduce pathological NOTCH3 accumulation while maintaining physiological NOTCH3 function. Alternatively, targeting downstream pathways regulated by NOTCH3 may provide another therapeutic avenue. Interestingly, rapamycin (sirolimus), an mTOR inhibitor, was identified through our proteomic analysis as a drug with similar predicted downstream effects to NOTCH3 knockdown. Rapamycin and its analogues are clinically used in drug-eluting stents to limit restenosis [102, 103] and function by inhibiting VSMC proliferation and migration [104, 105]. Furthermore, rapamycin has been shown to suppress the induction of CADASIL-associated proteins, including Decorin (DCN), Biglycan (BGN), and COL4A1, following exposure of primary human brain VSMCs to recombinant NOTCH3-Fc protein [35].

A limitation of this study is that the effects of the NOTCH3 siRNA were not assessed in a multicellular model. Therefore future studies will assess NOTCH3 siRNA in such models and further investigate whether combining NOTCH3 knockdown with replacement of functional NOTCH3 transcripts in CADASIL-derived mural cells can reduce pathological NOTCH3 accumulation while preserving physiological signalling. Collectively, these findings highlight that therapeutic modulation of NOTCH3 has the potential to influence multiple aspects of vascular cell biology and that the long-term consequences of sustained NOTCH3 suppression should be carefully evaluated during the development of RNA-based therapies for CADASIL and other NOTCH3-associated disorders.

In conclusion, this study demonstrates that NOTCH3 regulates ECM remodelling, cytoskeletal organisation, and glycolytic metabolism in human VSMCs while having limited effects proliferation, migration, and endothelial support. Importantly, characterisation of the NOTCH3 protein interactome identified a network of ECM-associated, cytoskeletal, and regulatory proteins providing new insights into the molecular mechanisms through which NOTCH3 contributes to VSMC homeostasis. Several molecular changes following NOTCH3 suppression were only identified following stimulation with PDGF or TGF-β, suggesting that NOTCH3 contributes to the regulation of VSMC responses to key environmental cues. These findings provide new insight into the physiological role of NOTCH3 in VSMC homeostasis and establish a framework for evaluating the potential consequences of therapeutic NOTCH3 suppression in CADASIL and other NOTCH3-related diseases. Together, these data support a model in which reduced physiological NOTCH3 signalling alters ECM production, cytoskeletal organisation, and metabolic pathways, particularly under conditions of vascular stimulation, highlighting NOTCH3 as an important regulator of VSMC state and adaptive responses.

## Abbreviations List

CADASIL: Cerebral Autosomal Dominant Arteriopathy with Subcortical Infarcts and Leukoencephalopathy
EC: Endothelial Cell
ECD: Extracellular Domain
EGFr: Epidermal Growth Factor-like repeats
DEPs: Differentially Expressed Proteins
GOM: Granular Osmiophilic Deposit
HAoSMC: Human Aortic Smooth Muscle Cell
ICD: Intracellular Domain
IPA: Ingenuity Pathway Analysis
PAH: Pulmonary Arterial Hypertension
PDGF: Platelet Derived Growth Factor-BB
SMC: Smooth Muscle Cell
TGF-β: Transforming Growth Factor-beta
VEGF: Vascular Endothelial Growth Factor
VSMC: Vascular Smooth Muscle Cell

## Author contributions

All experiments were performed in the Godson Lab in the Conway Institute at University College Dublin. **S.F.-** Conceptualization, design of work, data acquisition, data analysis, writing of manuscript, funding acquisition. **E.D.**-Data acquisition and analysis. **D.A.**-Methodology, interpretation of data. **K.J.B**.-Funding acquisition, interpretation of data, review of manuscript. **E.B.** -Resources, methodology, drafting of manuscript. **F.M.E.**-Methodology, design of work. **C.G.** -Supervision, design of work, resources, drafting of manuscript, funding acquisition.

## Acknowledgements

The authors would like to thank Mr Andrew Gaffney for his assistance at UCD Conway Labs and Professor Jeremy Simpson for providing access to the Opera Phenix Confocal Microscope. The authors acknowledge the use of ChatGPT (GPT-5.6, OpenAI, accessed July 2026) and Gemini (1.5 Pro, Google, accessed August 2026) solely for manuscript grammar refinement and code editing for the linear mixed-effects analyses.

## Funding

Horizon Europe Marie Skłodowska-Curie Actions (Call: HORIZON-MSCA-2022-PF-01, Project: CADASIL iMATTR, Grant ID: 101111254). UCD School of Medicine COINTREAU Fund (Grant ID: R28103). National Institute on Aging and Department of Veterans Affairs (IK2CX002180), Rainwater Charitable Foundation (Grant ID: 2790954), and Chan Zuckerberg Initiative (Grant ID: 2022-316712), and DataPhilanthropy (FME).

## Supplemental Figures

**Supplemental Table 1:**
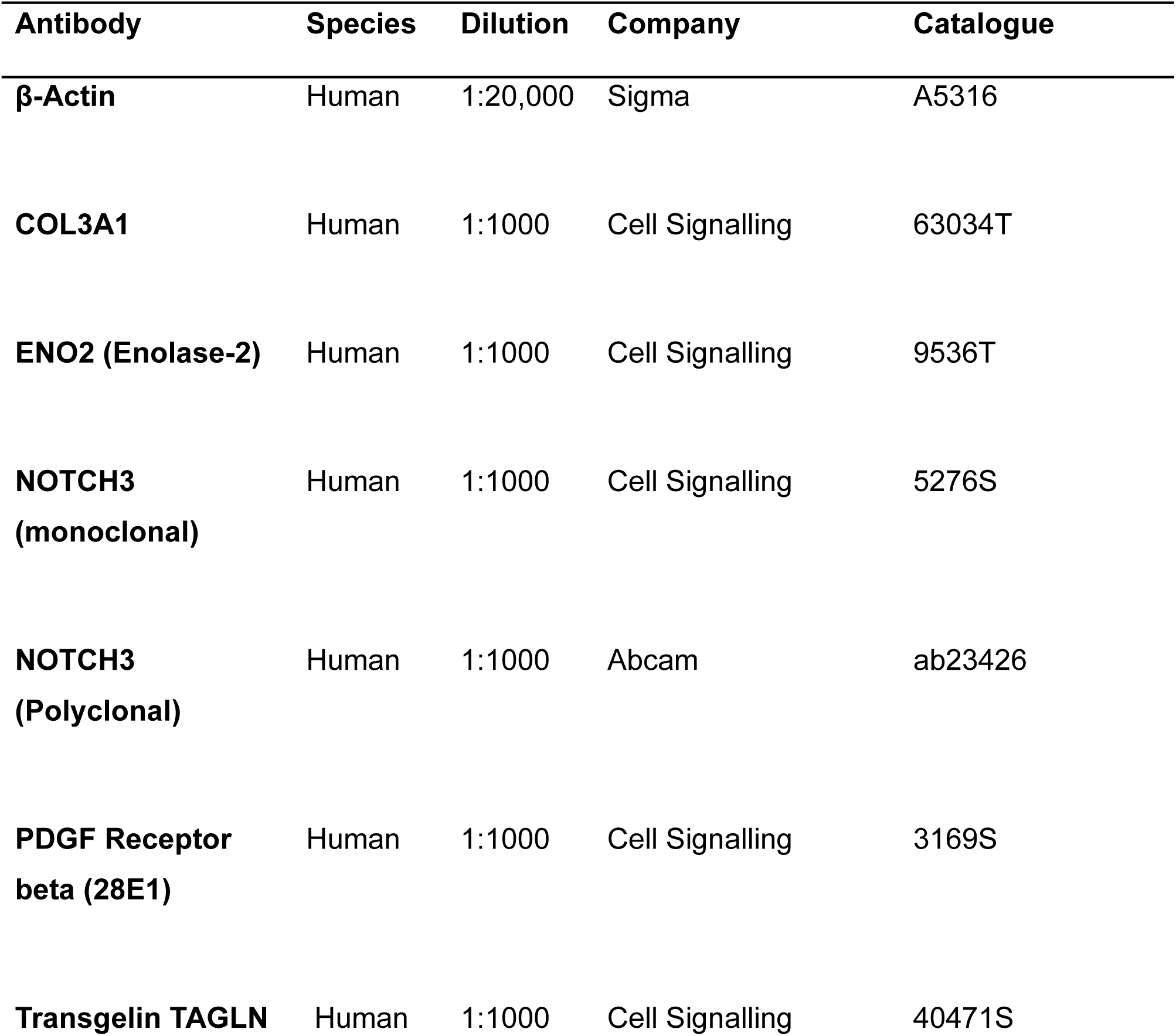
Western Blot Antibodies.

| **Antibody** | **Species** | **Dilution** | **Company** | **Catalogue** |
| --- | --- | --- | --- | --- |
| **β-Actin** | Human | 1:20,000 | Sigma | A5316 |
| **COL3A1** | Human | 1:1000 | Cell Signalling | 63034T |
| **ENO2 (Enolase-2)** | Human | 1:1000 | Cell Signalling | 9536T |
| **NOTCH3**<br>**(monoclonal)** | Human | 1:1000 | Cell Signalling | 5276S |
| **NOTCH3**<br>**(Polyclonal)** | Human | 1:1000 | Abcam | ab23426 |
| **PDGF Receptor**<br>**beta (28E1)** | Human | 1:1000 | Cell Signalling | 3169S |
| **Transgelin TAGLN** | Human | 1:1000 | Cell Signalling | 40471S |

**Supplemental Table 2:**
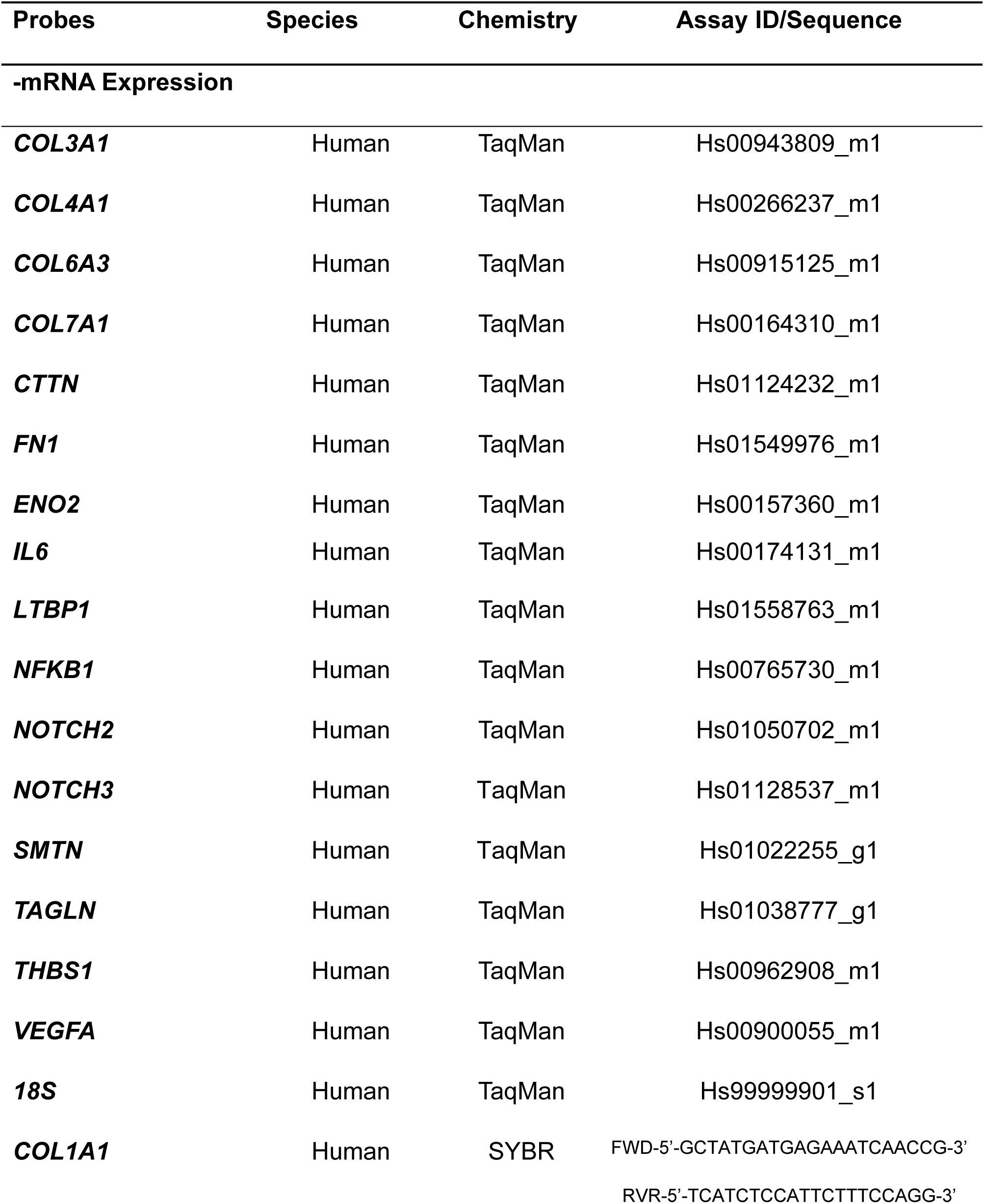
TaqMan and SYBR Assays used for qRT-PCR analysis.

**Supplemental Figure 1:**
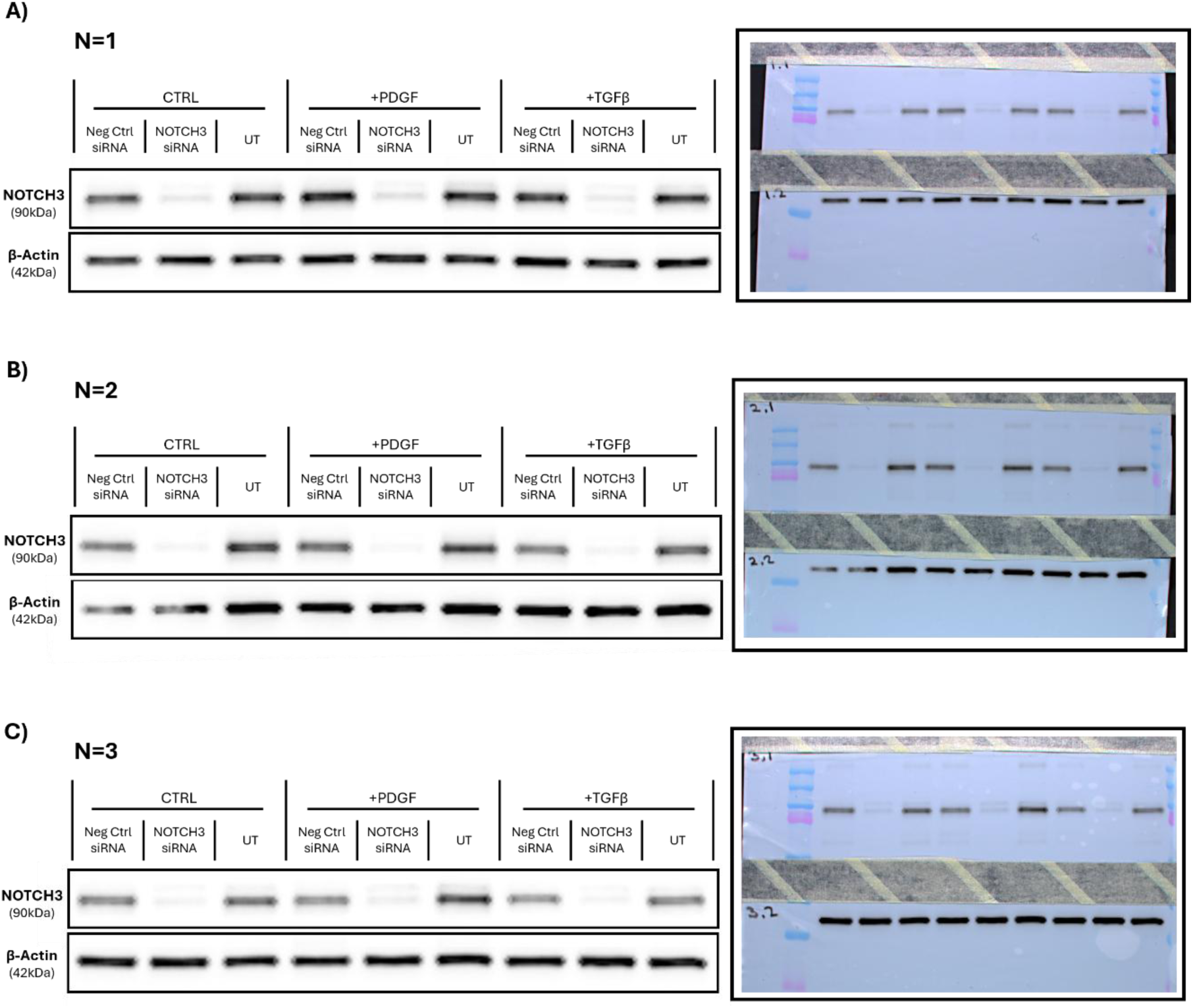
Raw blots of NOTCH3 following knockdown of NOTCH3 in HAoSMCs (N=3). HAoSMCs were treated with NOTCH3 siRNA or the negative control (Neg Ctrl) siRNA or left untransfected (UT) for 6 h followed by stimulation with PDGF (20 ng/mL), TGF-β (10 ng/mL) or vehicle control (CTRL; PBS with 0.05% BSA) for 48 h. NOTCH3 protein was analysed by Western blotting using β-actin as the control. **A-C)** Chemiluminescence signal intensity of labelled and cropped immunoblots, alongside representative images of the corresponding original blots from three independent experiments.

**Supplemental Figure 2:**
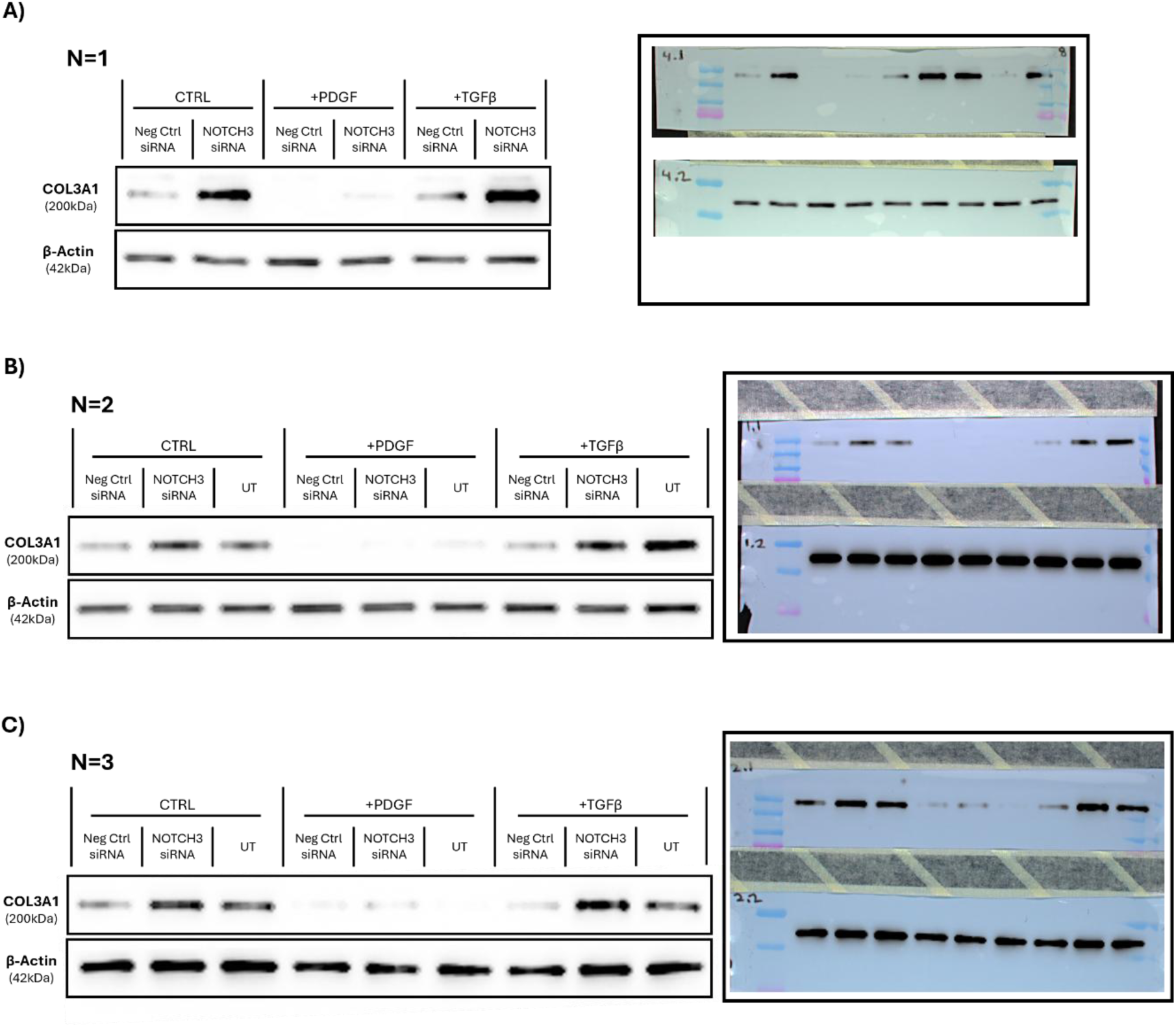
Raw blots of COL3A1 following knockdown of NOTCH3 in HAoSMCs (N=3). HAoSMCs were treated with NOTCH3 siRNA or the negative control (Neg Ctrl) siRNA or left untransfected (UT) for 6 h followed by stimulation with PDGF (20 ng/mL), TGF-β (10 ng/mL) or vehicle control (CTRL; PBS with 0.05% BSA) for 48 h. COL3A1 protein was analysed by Western blotting using β-actin as the control. **A-C)** Chemiluminescence signal intensity of labelled and cropped immunoblots, alongside representative images of the corresponding original blots from three independent experiments.

**Supplemental Figure 3:**
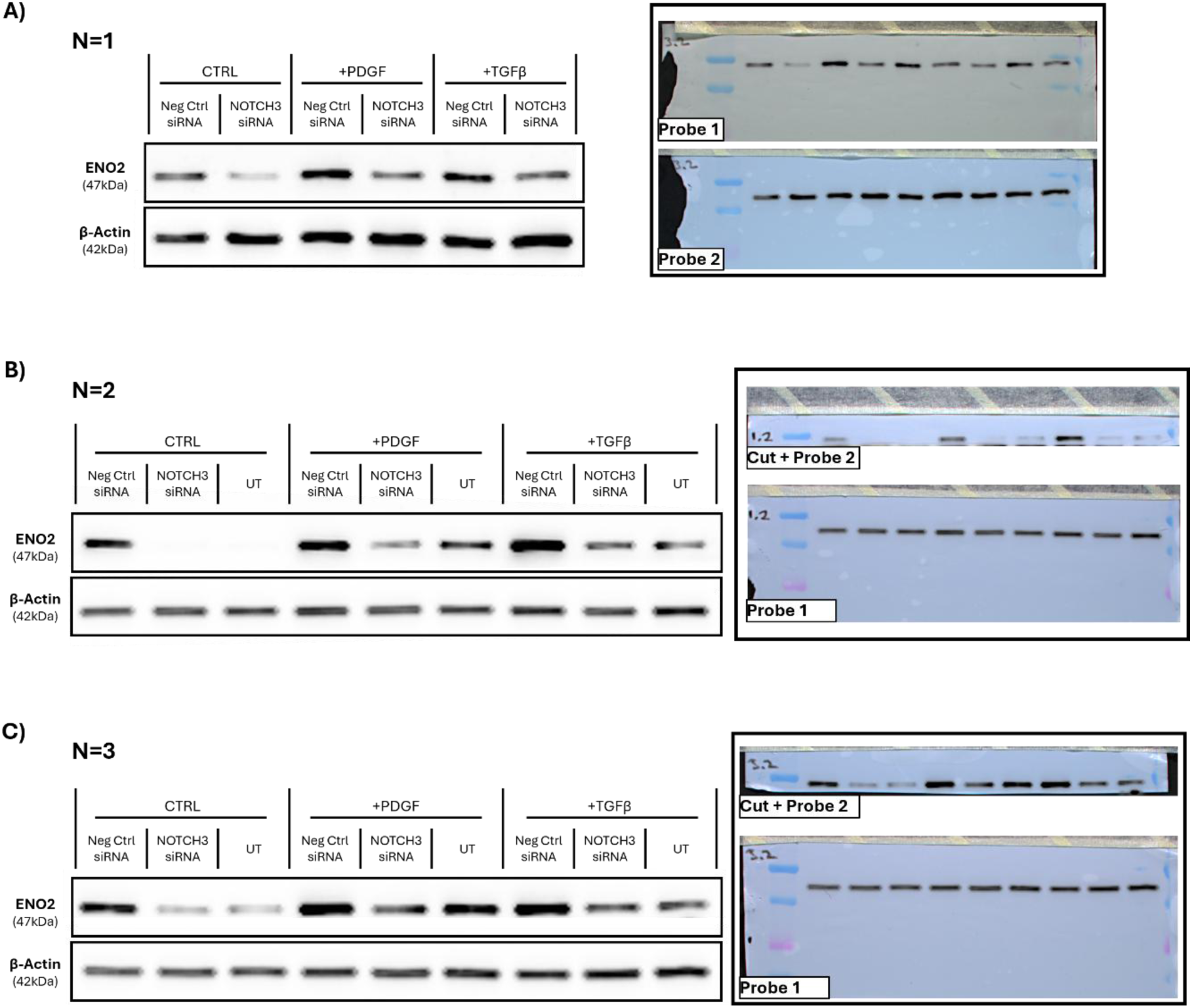
Raw blots of ENO2 following knockdown of NOTCH3 in HAoSMCs (N=3). HAoSMCs were treated with NOTCH3 siRNA or the negative control (Neg Ctrl) siRNA or left untransfected (UT) for 6 h followed by stimulation with PDGF (20 ng/mL), TGF-β (10 ng/mL) or vehicle control (CTRL; PBS with 0.05% BSA) for 48 h. ENO2 protein was analysed by Western blotting using β-actin as the control. **A-C)** Chemiluminescence signal intensity of labelled and cropped immunoblots, alongside representative images of the corresponding original blots from three independent experiments. Given the close proximity in molecular weight of ENO2 (47 kDa) to β-Actin (42kDa) immunoblots were cut and re-probed to prevent signal overlap.

**Supplemental Figure 4:**
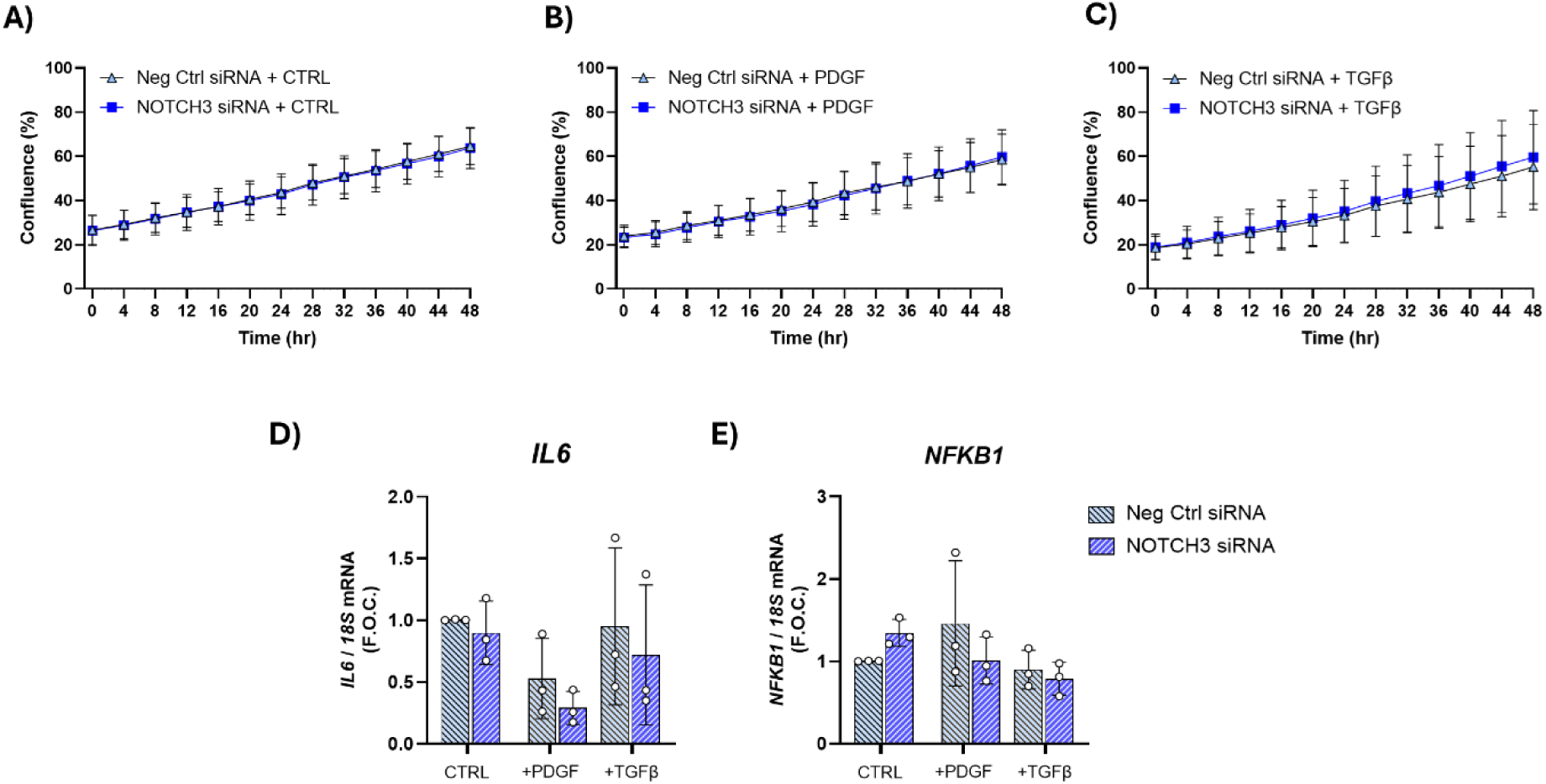
Effect of NOTCH3 siRNA on proliferation and inflammatory gene expression in HAoSMCs. HAoSMCs were treated with NOTCH3 siRNA or the negative control (Neg Ctrl) siRNA for 6 h followed by stimulation with PDGF (20 ng/mL), TGF-β (10 ng/mL) or vehicle control (CTRL; PBS with 0.05% BSA) for 48 h. **A-B)** Proliferation was analysed every 4 h using the Incucyte imaging system and quantified using the Incucyte software to calculate the percentage confluence across the three conditions (N=3,n=3). Gene expression analysis was performed using RT-PCR for **D)** *IL6* and **E)** *NFKB1* using 18S mRNA as the control (N=3). Statistical analysis was performed using an Ordinary One-Way ANOVA with Sidak’s multiple comparisons test.

**Supplemental Figure 5:**
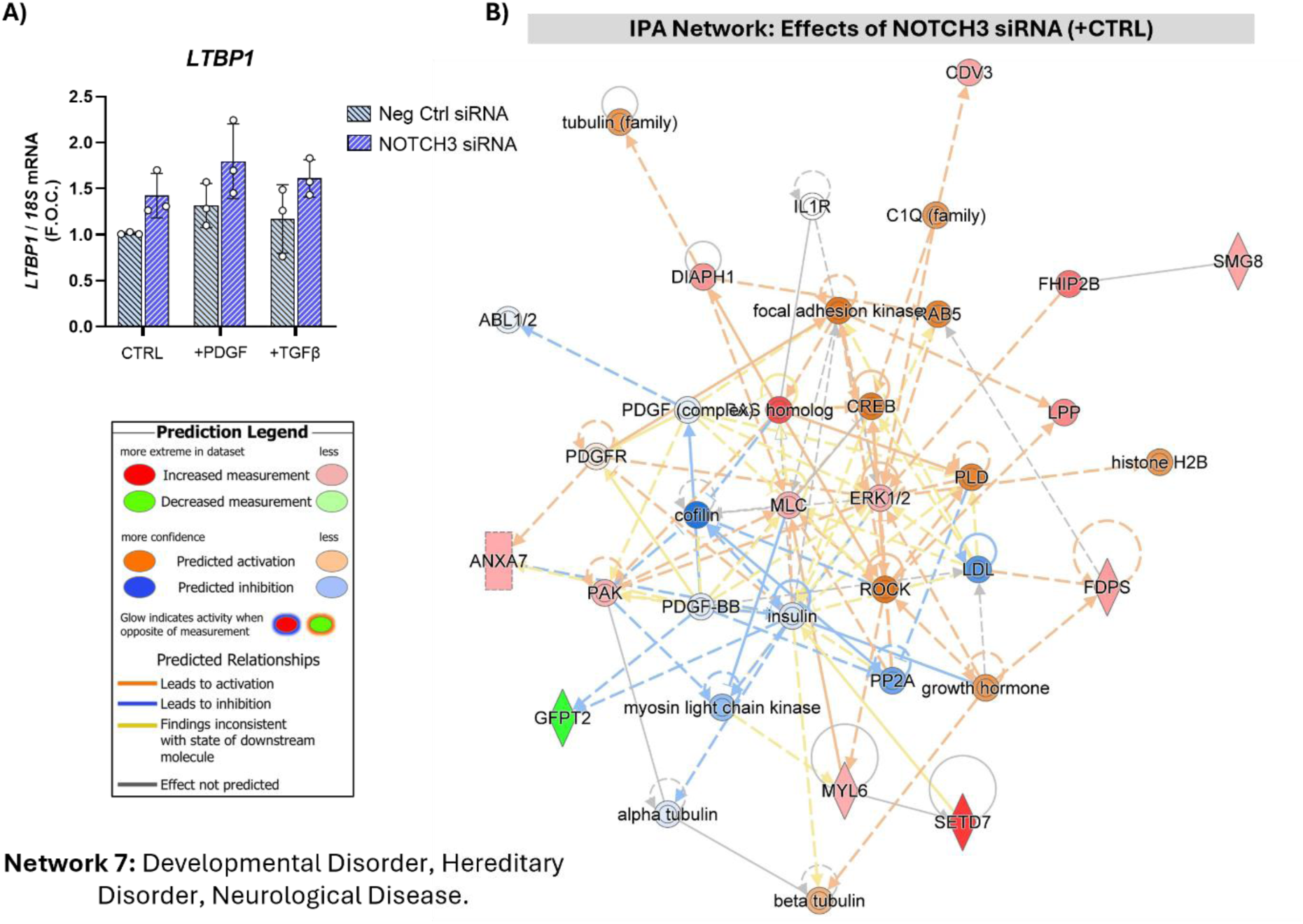
*LTBP1* expression and IPA analysis of the effects of NOTCH3 knockdown in HAoSMCs. HAoSMCs were treated with NOTCH3 siRNA or Neg Ctrl siRNA for 6 h followed by stimulation with PDGF (20 ng/mL), TGF-β (10 ng/mL) or vehicle control (CTRL; PBS with 0.05% BSA) for 48 h. **A)** Gene expression analysis of *LTBP1* by qRT-PCR using *18S rRNA* as the control (N=3). **B)** The list of differentially expressed proteins (DEPs) altered by NOTCH3 siRNA vs Neg Ctrl siRNA in the CTRL condition was analysed by IPA. SETD7 was identified as an interactor of MYL6 in Network 7 titled ‘Developmental Disorder, Hereditary Disorder, Neurological Disease’. Statistical analysis was performed using an Ordinary One-Way ANOVA with Sidak’s multiple comparisons test.

**Supplementary Figure 6:**
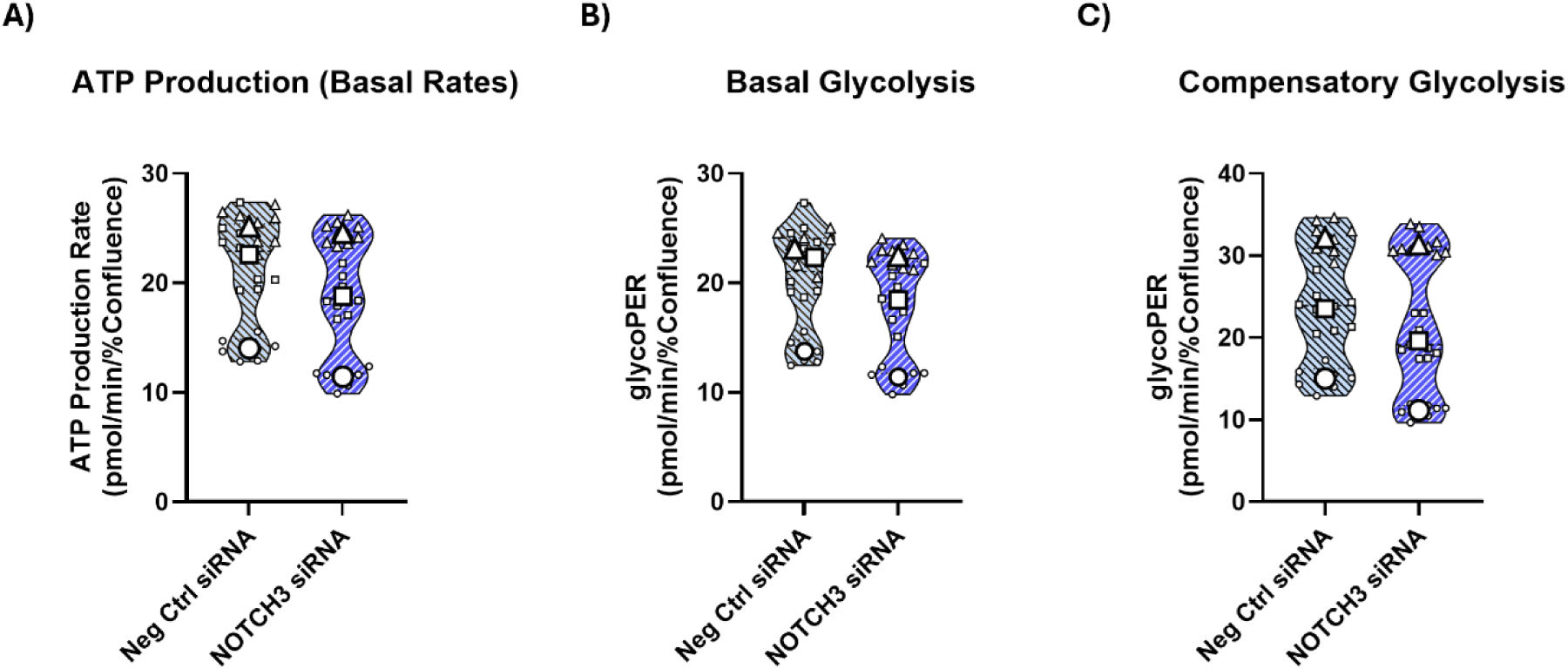
Analysis of ATP production, basal glycolysis and compensatory glycolysis in HAoSMCs following knockdown of NOTCH3. HAoSMCs were treated with NOTCH3 siRNA or Neg Ctrl siRNA for 6 h followed by a rest period of 24h. Cells were then seeded into a Seahorse XF96 Cell Culture Microplate and allowed to adhere overnight. **A)** ATP production, **B)** basal glycolysis, **C)** compensatory glycolysis were also assessed (N=3, n=8-9). Statistical analysis was performed on the mean values of the independent experiments using a paired T-test.

**Supplemental Figure 7:**
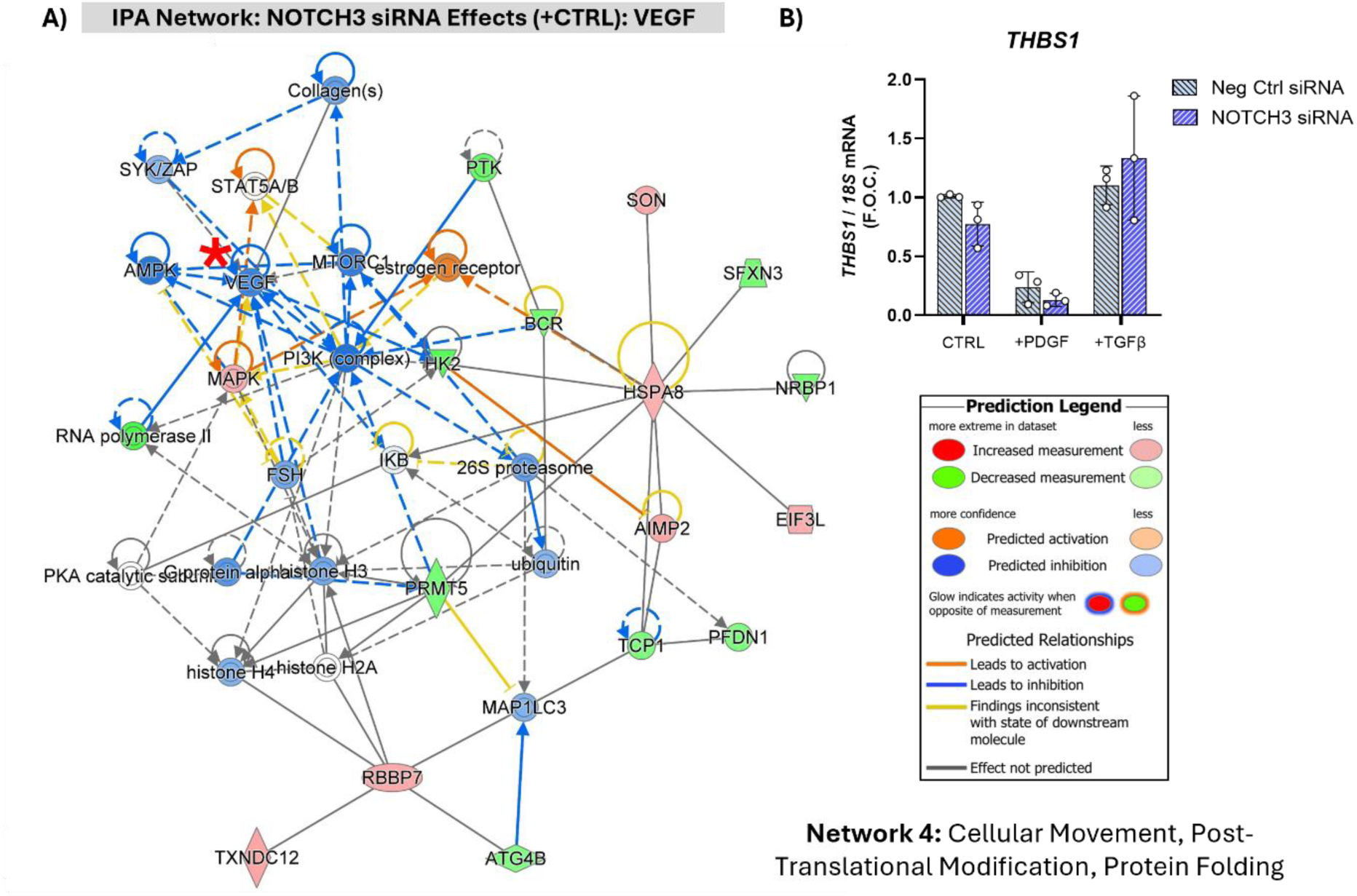
IPA analysis of DEPs and *THBS1* gene expression analysis of the effects of NOTCH3 knockdown in HAoSMCs. HAoSMCs were treated with NOTCH3 siRNA or the negative control (Neg Ctrl) siRNA for 6 h followed by stimulation with PDGF (20 ng/mL), TGF-β (10 ng/mL) or vehicle control (CTRL; PBS with 0.05% BSA) for 48 h. **A)** The list of differentially expressed proteins (DEPs) altered by NOTCH3 siRNA vs Negative control in the untreated (CTRL) condition was analysed by IPA. VEGF was identified as central node predicted to be downregulated (marked with a red asterisk (*)) in Network 4 titled ‘Cellular Movement, Post-Translational Modification, Protein Folding’. **B)** Gene expression analysis of *THBS1* by qRT-PCR using *18S rRNA* as the control (N=3). Statistical analysis was performed using an ordinary one-way ANOVA with Sidak’s multiple comparisons test.

**Supplemental Figure 8:**
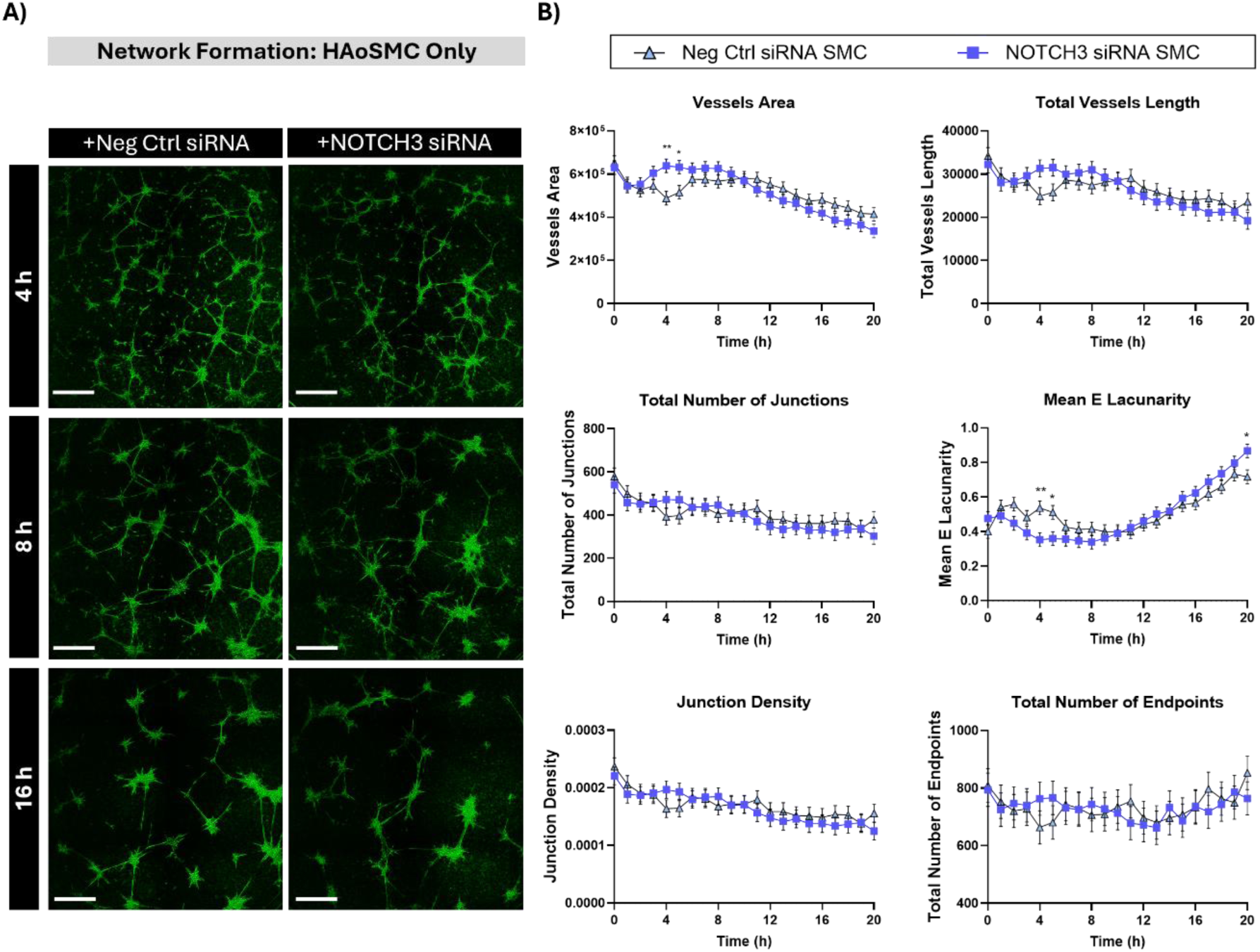
Knockdown of NOTCH3 in HAoSMCs has minimal effects on the HAoSMC interaction network. HAoSMCs were treated with NOTCH3 siRNA for 6 h using Neg Ctrl siRNA as the control. Cells were rested and then seeded on Geltrex and analysed by the Incucyte for 20 h. **A)** Representative images of SMC networks after 4, 8 and 16 h across the two conditions. Brightfield images were pseudocoloured green for visualisation purposes. Scale bar = 800 µm. **B)** Quantitative analysis of the network across all time points was performed using AngioTool2.0 Software (N=4, n=3). A linear mixed effects model was fitted to the data and values depict the estimated marginal means ± SE (N=4, n=3). Statistical comparisons between groups were performed at each time point using estimated marginal means with Holm correction for multiple comparisons. *=p<0.05, **=p<0.01.

**Supplemental Figure 9:**
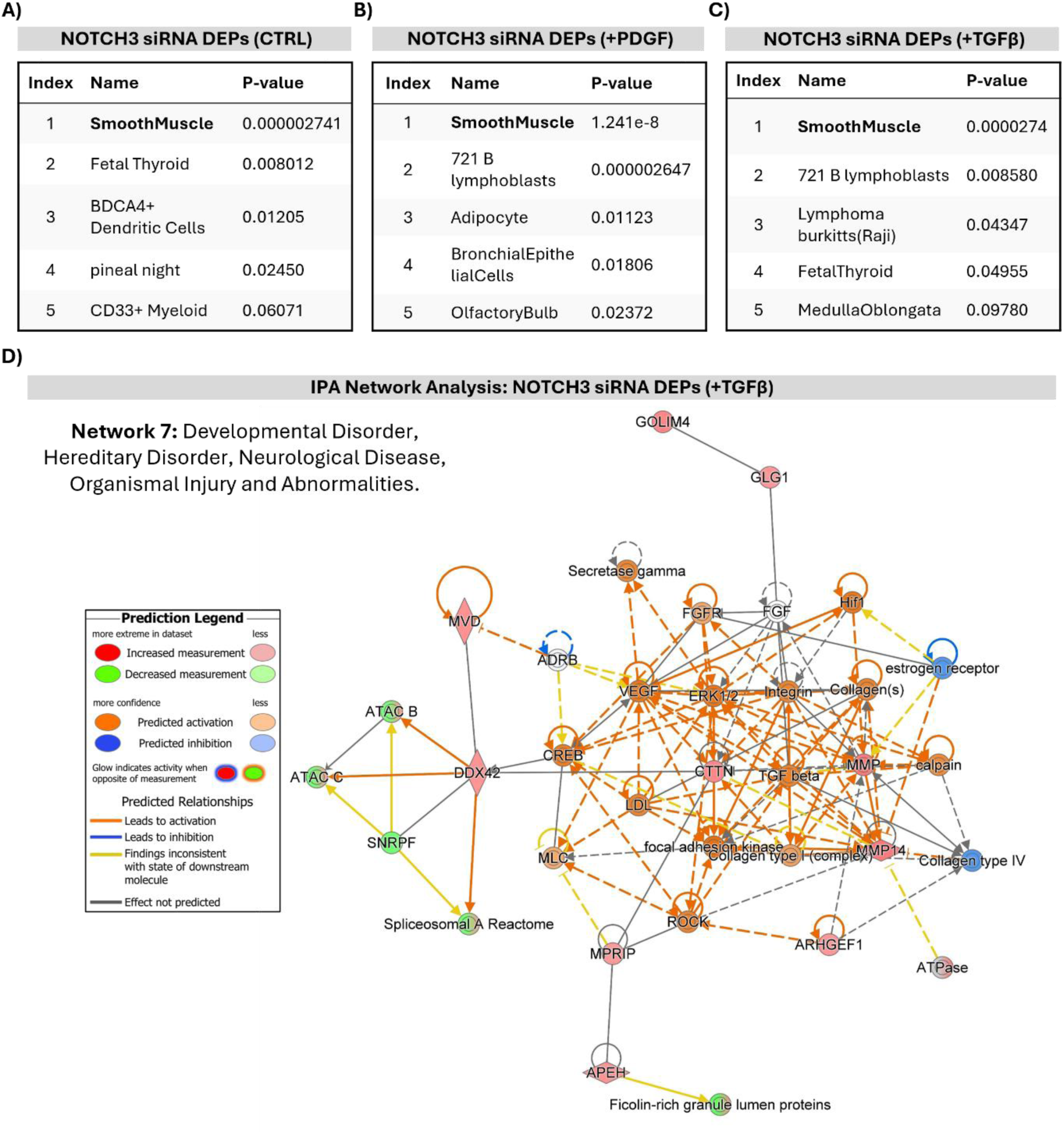
ENRICHR and IPA analysis of differentially expressed proteins in HAoSMCs following knockdown of NOTCH3. HAoSMCs were treated with NOTCH3 siRNA or the negative control (Neg Ctrl) siRNA or left untransfected (UT) for 6 h followed by stimulation with PDGF (20 ng/mL), TGF-β (10 ng/mL) or vehicle control (CTRL; PBS with 0.05% BSA) for 48 h. **A-C)** Proteomic analysis was performed and the differentially expressed proteins (DEPs) in each condition were analysed by the Human Gene Atlas via ENRICHR Software. **D)** The list of DEPs altered by NOTCH3 siRNA in the presence of TGFβ was analysed by IPA. CTTN was increased and collagens and TGFβ were predicted to be activated in IPA Network 7 titled ‘Developmental Disorder, Hereditary Disorder, Neurological Disease, Organismal Injury and Abnormalities’.

**Supplemental Figure 10:**
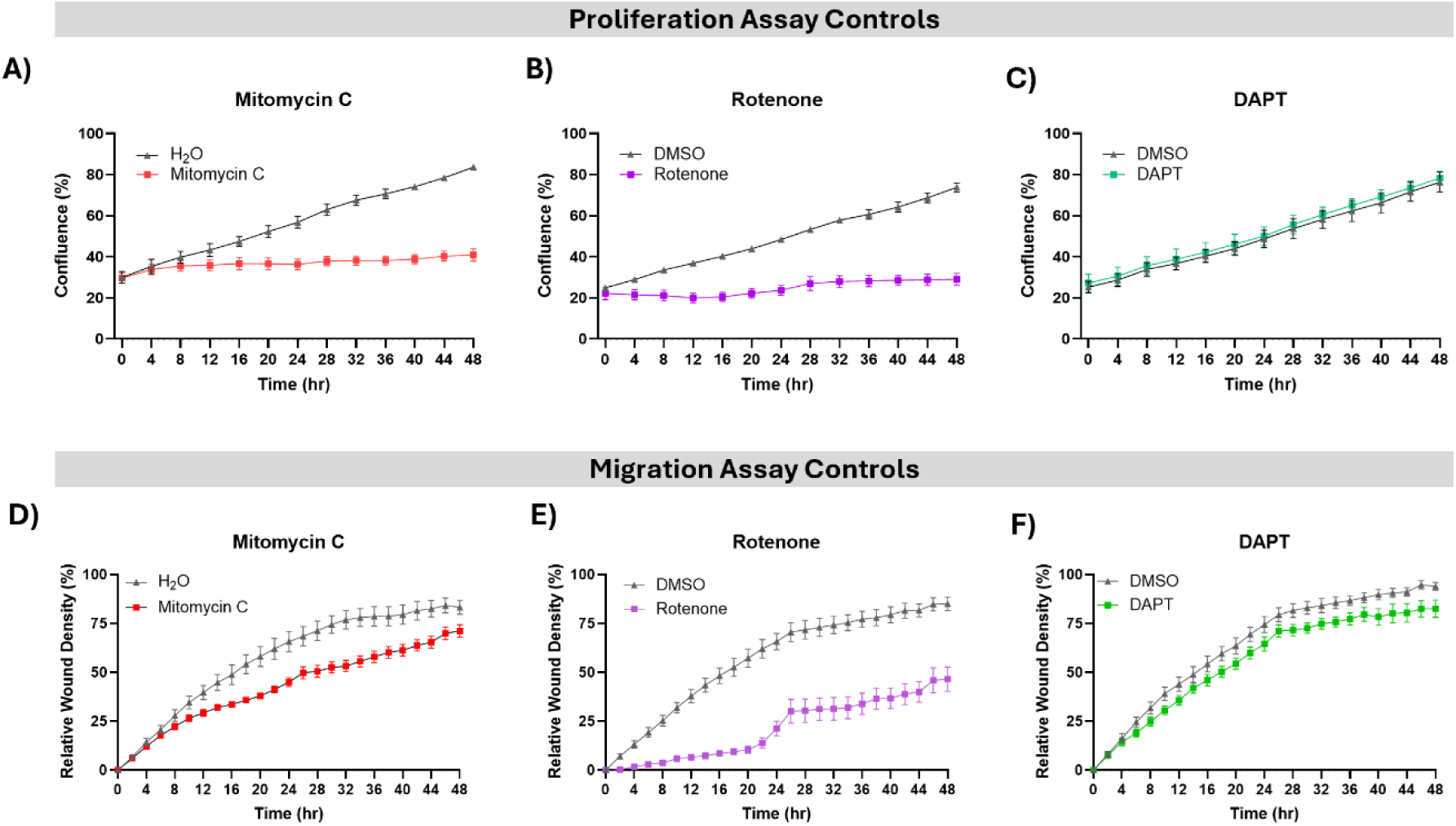
HAoSMC proliferation and migration assay controls. HAoSMCs were treated with NOTCH3 siRNA or the negative control (Neg Ctrl) siRNA or left untransfected (UT) for 6 h followed by stimulation with PDGF (20 ng/mL), TGF-β (10 ng/mL) or vehicle control (CTRL; PBS with 0.05% BSA) for 48 h. On the same plate HAoSMCs were seeded but left untransfected. Following overnight rest, these untransfected cells were treated for 48 h with the assay controls **A)** Mitomycin C (20 µg/mL) using H20 as the vehicle control; **B)** Rotenone (25 µM) using DMSO as the vehicle control; **C)** the γ-secretase inhibitor, DAPT, using DMSO as the control. Live cell proliferation analysis was initiated after treatment (0 h) for a total of 48 h using the Incucyte. Separately, HAoSMCs were seeded and treated with NOTCH3 siRNA for 6hr for the migration assay alongside untreated cells. After overnight rest, all cells were incubated with Mitomycin-C (20 µg/mL) for 2 h to inhibit proliferation. A scratch was generated using the WoundMaker from Incuycte and the untransfected cells were stimulated for 48 h with the assay controls **D)** Mitomycin-C (20 µg/mL) using H20 as the vehicle control; **E)** Rotenone (25 µM) using DMSO as the vehicle control; **F)** the γ-secretase inhibitor, DAPT, using DMSO as the control. Cells were analysed every 4 h for a total of 48 h. The images were obtained using Incucyte live cell imaging every 4 h for a total of 48 h. Each independent experiment consisted of three biological replicates analysed per condition. Error bars are representative of the average of 3 independent experiments.

**Supplemental Figure 11:**
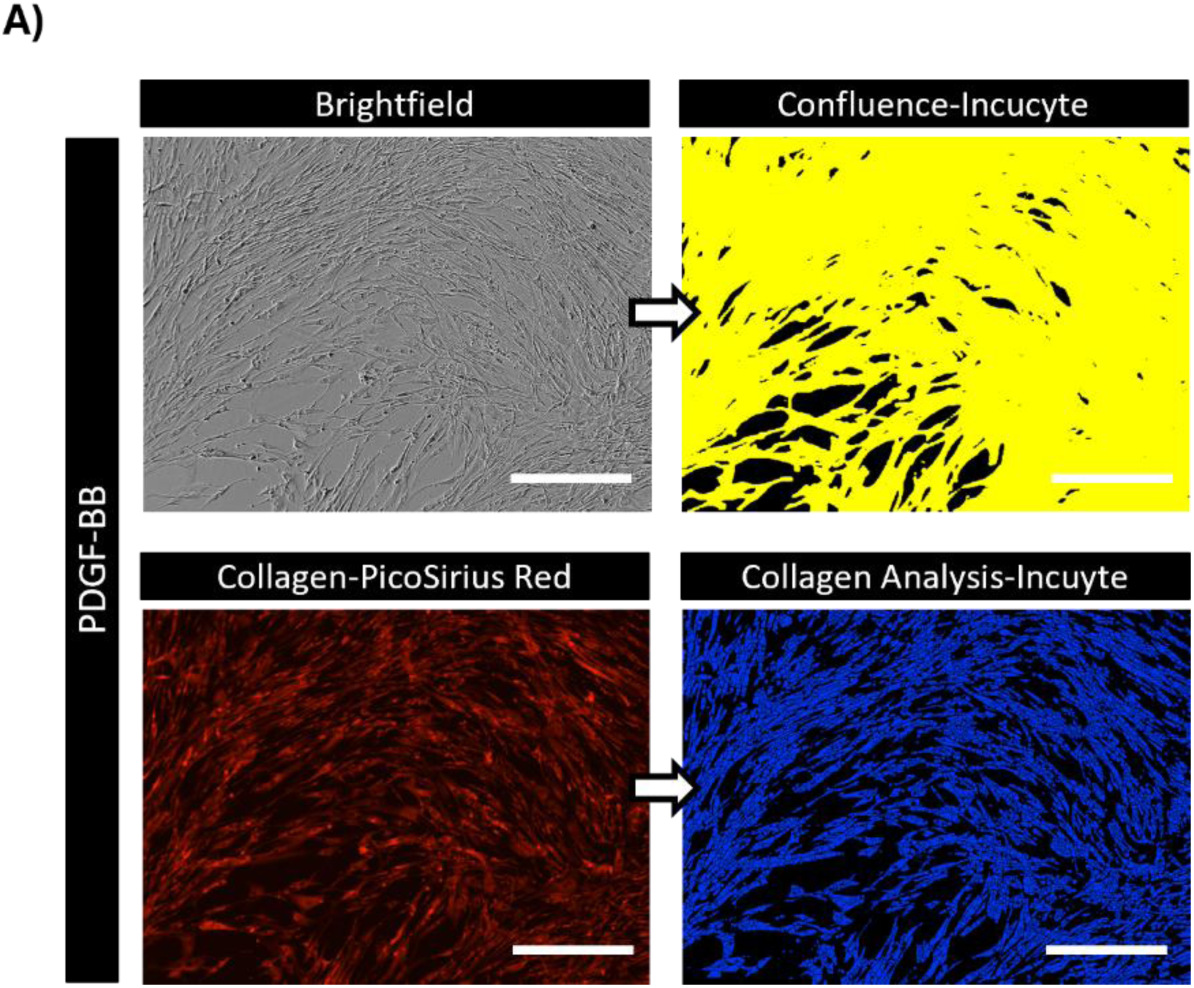
Collagen staining assessment and cell confluence assessment via Incucyte. HAoSMCs were treated with NOTCH3 siRNA or Neg Ctrl siRNA for 6 h followed by stimulation with PDGF-BB (20 ng/mL) and TGF-β (10 ng/mL) or vehicle control (CTRL; PBS with 0.01% BSA) for 4 days. **A)** The Incucyte was used to obtain representative images of the cells by brightfield and by fluorescence for PicoSirius Red. Incuycte software was used to quantify the confluence (yellow overlay) and the red fluorescence intensity (blue overlay). Scale bar = 400 µm.

**Supplemental Figure 12:**
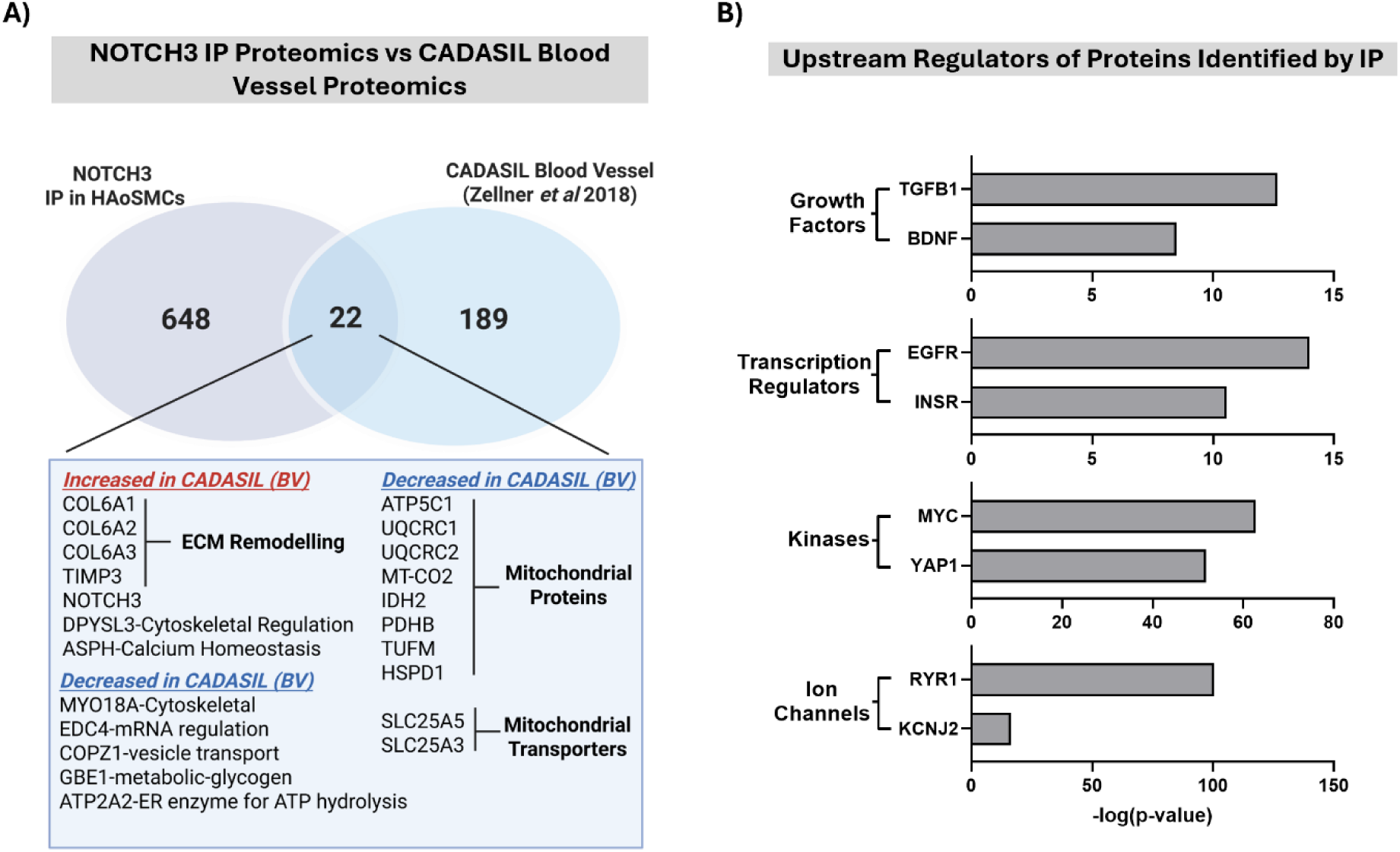
**A)** Intersection of NOTCH3 interactome as identified by NOTCH3 immunoprecipitation with previously published proteomic dataset analysis of brain-derived CADASIL blood vessel [26]. **B)** Top upstream regulators predicted to be activated by NOTCH3 interactome identified by IPA (Qiagen) categorised by ‘Molecule Type’ and graphed based on statistical significance.

## Notes

### Competing Interest Statement

The authors have declared no competing interest.

## References

1. Armulik, A., A. Abramsson, and C. Betsholtz, Endothelial/Pericyte Interactions. Circulation Research, 2005. 97(6): p. 512–523.

2. Rensen, S.S., P.A. Doevendans, and G.J. van Eys, Regulation and characteristics of vascular smooth muscle cell phenotypic diversity. Neth Heart J, 2007. 15(3): p. 100–8.

3. Owens, G.K., M.S. Kumar, and B.R. Wamhoff, Molecular Regulation of Vascular Smooth Muscle Cell Differentiation in Development and Disease. Physiological Reviews, 2004. 84(3): p. 767–801.

4. Natali, L., et al., A Comprehensive Review of Epigenetic Regulation of Vascular Smooth Muscle Cells During Development and Disease. Biomolecules, 2026. 16(1).

5. Lambert, J. and H.F. Jørgensen, Epigenetic regulation of vascular smooth muscle cell phenotypes in atherosclerosis. Atherosclerosis, 2025. 401: p. 119085.

6. Chakraborty, R., et al., Targeting smooth muscle cell phenotypic switching in vascular disease. JVS Vasc Sci, 2021. 2: p. 79–94.

7. Domenga, V., et al., Notch3 is required for arterial identity and maturation of vascular smooth muscle cells. Genes Dev, 2004. 18(22): p. 2730–5.

8. Liu, H., et al., Notch3 is critical for proper angiogenesis and mural cell investment. Circ Res, 2010. 107(7): p. 860–70.

9. Del Gaudio, F., D. Liu, and U. Lendahl, Notch signalling in healthy and diseased vasculature. Open Biology, 2022. 12(4).

10. Bodas, M., et al., The NOTCH3 Downstream Target HEYL Is Required for Efficient Human Airway Basal Cell Differentiation. Cells, 2021. 10(11).

11. Liu, H., S. Kennard, and B. Lilly, NOTCH3 Expression Is Induced in Mural Cells Through an Autoregulatory Loop That Requires Endothelial-Expressed JAGGED1. Circulation Research, 2009. 104(4): p. 466–475.

12. Hoglund, V.J. and M.W. Majesky, Patterning the Artery Wall by Lateral Induction of Notch Signaling. Circulation, 2012. 125(2): p. 212–215.

13. Regan, J.N. and M.W. Majesky, Building a vessel wall with notch signaling. Circ Res, 2009. 104(4): p. 419–21.

14. Manderfield, L.J., et al., Notch activation of Jagged1 contributes to the assembly of the arterial wall. Circulation, 2012. 125(2): p. 314–23.

15. Liu, H., S. Kennard, and B. Lilly, NOTCH3 expression is induced in mural cells through an autoregulatory loop that requires endothelial-expressed JAGGED1. Circ Res, 2009. 104(4): p. 466–75.

16. Romay, M.C., et al., Age-related loss of Notch3 underlies brain vascular contractility deficiencies, glymphatic dysfunction, and neurodegeneration in mice. The Journal of Clinical Investigation, 2024. 134(2).

17. Li, X., et al., Notch3 signaling promotes the development of pulmonary arterial hypertension. Nature Medicine, 2009. 15(11): p. 1289–1297.

18. Hernandez, M., et al., The NOTCH3 extracellular domain is a serum biomarker for pulmonary arterial hypertension. Nature Medicine, 2026. 32(1): p. 306–317.

19. Aburjania, Z., et al., The Role of Notch3 in Cancer. Oncologist, 2018. 23(8): p. 900–911.

20. Kondratyev, M., et al., Identification of acquired Notch3 dependency in metastatic Head and Neck Cancer. Communications Biology, 2023. 6(1): p. 538.

21. Xiang, H., et al., Single-Cell Analysis Identifies NOTCH3-Mediated Interactions between Stromal Cells That Promote Microenvironment Remodeling and Invasion in Lung Adenocarcinoma. Cancer Res, 2024. 84(9): p. 1410–1425.

22. Zhang, Z., et al., Notch3 in human breast cancer cell lines regulates osteoblast-cancer cell interactions and osteolytic bone metastasis. Am J Pathol, 2010. 177(3): p. 1459–69.

23. Lee, S.J., et al., Structural changes in NOTCH3 induced by CADASIL mutations: Role of cysteine and non-cysteine alterations. J Biol Chem, 2023. 299(6): p. 104838.

24. Gatti, J.R., et al., Redistribution of Mature Smooth Muscle Markers in Brain Arteries in Cerebral Autosomal Dominant Arteriopathy with Subcortical Infarcts and Leukoencephalopathy. Transl Stroke Res, 2018.

25. Dong, H., M. Blaivas, and M.M. Wang, Bidirectional encroachment of collagen into the tunica media in cerebral autosomal dominant arteriopathy with subcortical infarcts and leukoencephalopathy. Brain Research, 2012. 1456: p. 64–71.

26. Zellner, A., et al., CADASIL brain vessels show a HTRA1 loss-of-function profile. Acta Neuropathol, 2018. 136(1): p. 111–125.

27. Tikka, S., et al., CADASIL mutations and shRNA silencing of NOTCH3 affect actin organization in cultured vascular smooth muscle cells. J Cereb Blood Flow Metab, 2012. 32(12): p. 2171–80.

28. Joutel, A., et al., The ectodomain of the Notch3 receptor accumulates within the cerebrovasculature of CADASIL patients. The Journal of clinical investigation, 2000. 105(5): p. 597–605.

29. Hack, R.J., et al., Three-tiered EGFr domain risk stratification for individualized NOTCH3-small vessel disease prediction. Brain, 2023. 146(7): p. 2913–2927.

30. Chabriat, H., et al., Cadasil. Lancet Neurol, 2009. 8(7): p. 643–53.

31. Jolly, A.A., et al., Prevalence of Fatigue and Associations With Depression and Cognitive Impairment in Patients With CADASIL. Neurology, 2025. 104(3): p. e213335.

32. Karvelas, N., et al., Defining patient-reported outcomes and priorities for clinical trials in CADASIL through an international survey. Cerebral Circulation -Cognition and Behavior, 2026. 10: p. 100534.

33. Dupré, N., et al., Protein aggregates containing wild-type and mutant NOTCH3 are major drivers of arterial pathology in CADASIL. The Journal of Clinical Investigation, 2024. 134(8).

34. Mizuta, I., et al., Progress to Clarify How NOTCH3 Mutations Lead to CADASIL, a Hereditary Cerebral Small Vessel Disease. Biomolecules, 2024. 14(1).

35. Zhang, X., et al., The small leucine-rich proteoglycan BGN accumulates in CADASIL and binds to NOTCH3. Transl Stroke Res, 2015. 6(2): p. 148–55.

36. Monet-Leprêtre, M., et al., Abnormal recruitment of extracellular matrix proteins by excess Notch3 ECD: a new pathomechanism in CADASIL. Brain, 2013. 136(Pt 6): p. 1830–45.

37. Donahue, C.P. and K.S. Kosik, Distribution pattern of Notch3 mutations suggests a gain-of-function mechanism for CADASIL. Genomics, 2004. 83(1): p. 59–65.

38. Long, L., et al., Reduced SUMOylation impairs NOTCH3 signaling and cell survival in the pathogenesis of CADASIL. Cell Commun Signal, 2025. 24(1): p. 3.

39. Ling, C., et al., Modeling CADASIL vascular pathologies with patient-derived induced pluripotent stem cells. Protein & Cell, 2019. 10(4): p. 249–271.

40. Pippucci, T., et al., Homozygous NOTCH3 null mutation and impaired NOTCH3 signaling in recessive early-onset arteriopathy and cavitating leukoencephalopathy. EMBO Mol Med, 2015. 7(6): p. 848–58.

41. Watanabe-Hosomi, A., et al., Transendocytosis is impaired in CADASIL-mutant NOTCH3. Exp Neurol, 2012. 233(1): p. 303–11.

42. Suzuki, S., et al., Lunatic fringe promotes the aggregation of CADASIL NOTCH3 mutant proteins. Biochemical and Biophysical Research Communications, 2021. 557: p. 302–308.

43. Kelleher, J., et al., Patient-Specific iPSC Model of a Genetic Vascular Dementia Syndrome Reveals Failure of Mural Cells to Stabilize Capillary Structures. Stem Cell Reports, 2019. 13(5): p. 817–831.

44. Gastfriend, B.D., et al., Notch3 directs differentiation of brain mural cells from human pluripotent stem cell–derived neural crest. Science Advances, 2024. 10(5): p. eadi1737.

45. Zaucker, A., et al., notch3 is essential for oligodendrocyte development and vascular integrity in zebrafish. Dis Model Mech, 2013. 6(5): p. 1246–59.

46. Ando, K., et al., Peri-arterial specification of vascular mural cells from naïve mesenchyme requires Notch signaling. Development, 2019. 146(2).

47. Henshall, T.L., et al., Notch3 is necessary for blood vessel integrity in the central nervous system. Arterioscler Thromb Vasc Biol, 2015. 35(2): p. 409–20.

48. Haile, S., et al., Cerebral autosomal dominant arteriopathy with subcortical infarcts and leukoencephalopathy (CADASIL): potential therapeutic approaches. The Nucleus, 2025. 68(3): p. 471–487.

49. Rutten, J.W., et al., Therapeutic NOTCH3 cysteine correction in CADASIL using exon skipping: in vitro proof of concept. Brain, 2016. 139(Pt 4): p. 1123–35.

50. Livak, K.J. and T.D. Schmittgen, Analysis of Relative Gene Expression Data Using Real-Time Quantitative PCR and the 2−ΔΔCT Method. Methods, 2001. 25(4): p. 402–408.

51. Fitzsimons, S., et al., Inhibition of pro-inflammatory signaling in human primary macrophages by enhancing arginase-2 via target site blockers. Mol Ther Nucleic Acids, 2023. 33: p. 941–959.

52. Perez-Riverol, Y., et al., The PRIDE database and related tools and resources in 2019: improving support for quantification data. Nucleic Acids Res, 2019. 47(D1): p. D442–d450.

53. Turriziani, B., et al., On-Beads Digestion in Conjunction with Data-Dependent Mass Spectrometry: A Shortcut to Quantitative and Dynamic Interaction Proteomics. Biology, 2014. 3(2): p. 320–332.

54. Rappsilber, J., M. Mann, and Y. Ishihama, Protocol for micro-purification, enrichment, pre-fractionation and storage of peptides for proteomics using StageTips. Nature protocols, 2007. 2(8): p. 1896–1906.

55. Baeten, J.T. and B. Lilly, Differential Regulation of NOTCH2 and NOTCH3 Contribute to Their Unique Functions in Vascular Smooth Muscle Cells. J Biol Chem, 2015. 290(26): p. 16226–37.

56. Lindahl, P., et al., Pericyte Loss and Microaneurysm Formation in PDGF-B-Deficient Mice. Science, 1997. 277(5323): p. 242–245.

57. Bjarnegård, M., et al., Endothelium-specific ablation of PDGFB leads to pericyte loss and glomerular, cardiac and placental abnormalities. Development, 2004. 131(8): p. 1847–1857.

58. Martin-Garrido, A., et al., NADPH oxidase 4 mediates TGF-β-induced smooth muscle α-actin via p38MAPK and serum response factor. Free Radical Biology and Medicine, 2011. 50(2): p. 354–362.

59. Verrecchia, F., M.-L. Chu, and A. Mauviel, Identification of Novel TGF-&#x3b2;/Smad Gene Targets in Dermal Fibroblasts using a Combined cDNA Microarray/Promoter Transactivation Approach *. Journal of Biological Chemistry, 2001. 276(20): p. 17058–17062.

60. Zhang, Y.-Q., et al., Notch3 inhibits cell proliferation and tumorigenesis and predicts better prognosis in breast cancer through transactivating PTEN. Cell Death & Disease, 2021. 12(6): p. 502.

61. Kuivaniemi, H. and G. Tromp, Type III collagen (COL3A1): Gene and protein structure, tissue distribution, and associated diseases. Gene, 2019. 707: p. 151–171.

62. Singh, D., V. Rai, and D.K. Agrawal, Regulation of Collagen I and Collagen III in Tissue Injury and Regeneration. Cardiol Cardiovasc Med, 2023. 7(1): p. 5–16.

63. Kast, J., et al., Sequestration of latent TGF-β binding protein 1 into CADASIL-related Notch3-ECD deposits. Acta Neuropathol Commun, 2014. 2: p. 96.

64. Lin, H., et al., Annexin A2 promotes angiogenesis after ischemic stroke via annexin A2 receptor – AKT/ERK pathways. Neuroscience Letters, 2023. 792: p. 136941.

65. Sjöberg, E., et al., Endothelial VEGFR2-PLCγ signaling regulates vascular permeability and antitumor immunity through eNOS/Src. The Journal of Clinical Investigation, 2023. 133(20).

66. Martin, K.A., et al., The mTOR/p70 S6K1 pathway regulates vascular smooth muscle cell differentiation. American Journal of Physiology-Cell Physiology, 2004. 286(3): p. C507–C517.

67. Martin, K.A., et al., Rapamycin Promotes Vascular Smooth Muscle Cell Differentiation through Insulin Receptor Substrate-1/Phosphatidylinositol 3-Kinase/Akt2 Feedback Signaling *. Journal of Biological Chemistry, 2007. 282(49): p. 36112–36120.

68. Shuttleworth, V.G., et al., The methyltransferase SETS regulates TGFB1 activation of renal fibroblasts via interaction with SMAD3. J Cell Sci, 2018. 131(1).

69. Mohammed, S.A., et al., Targeting SETD7 Rescues Diabetes-Induced Impairment of Angiogenic Response by Transcriptional Repression of Semaphorin-3G. Diabetes, 2025. 74(6): p. 969–982.

70. Elkouris, M., et al., SET9-Mediated Regulation of TGF-β Signaling Links Protein Methylation to Pulmonary Fibrosis. Cell Reports, 2016. 15(12): p. 2733–2744.

71. Viitanen, M., et al., Experimental studies of mitochondrial function in CADASIL vascular smooth muscle cells. Experimental Cell Research, 2013. 319(3): p. 134–143.

72. Annunen-Rasila, J., et al., Cytoskeletal structure in cells harboring two mutations: R133C in NOTCH3 and 5650G>A in mitochondrial DNA. Mitochondrion, 2007. 7(1-2): p. 96–100.

73. Schröder, J.M., et al., Peripheral nerve and skeletal muscle involvement in CADASIL. Acta Neuropathol, 2005. 110(6): p. 587–99.

74. Zheng, Y., et al., Insulin-like growth factor 1-induced enolase 2 deacetylation by HDAC3 promotes metastasis of pancreatic cancer. Signal Transduction and Targeted Therapy, 2020. 5(1): p. 53.

75. Zhao, X., et al., Selective vulnerability of cerebral vasculature to <em>NOTCH3</em> variants in small vessel disease and rescue by phosphodiesterase-5 inhibitor. bioRxiv, 2025: p. 2025.05.30.656914.

76. Weaver, A.M., et al., Cortactin promotes and stabilizes Arp2/3-induced actin filament network formation. Current Biology, 2001. 11(5): p. 370–374.

77. Zhang, X., H. Meng, and M.M. Wang, Collagen represses canonical Notch signaling and binds to Notch ectodomain. Int J Biochem Cell Biol, 2013. 45(7): p. 1274–80.

78. Wu, L., et al., Identification of a family of mastermind-like transcriptional coactivators for mammalian notch receptors. Mol Cell Biol, 2002. 22(21): p. 7688–700.

79. Beres, B.J., et al., Numb regulates Notch1, but not Notch3, during myogenesis. Mechanisms of Development, 2011. 128(5): p. 247–257.

80. Dabertrand, F., et al., PIP 2 corrects cerebral blood flow deficits in small vessel disease by rescuing capillary Kir2.1 activity. Proceedings of the National Academy of Sciences, 2021. 118(17): p. e2025998118.

81. Huneau, C., et al., Altered dynamics of neurovascular coupling in CADASIL. Ann Clin Transl Neurol, 2018. 5(7): p. 788–802.

82. Tikka, S., et al., CADASIL and CARASIL. Brain Pathology, 2014. 24(5): p. 525–544.

83. Dong, H., M. Blaivas, and M.M. Wang, Bidirectional encroachment of collagen into the tunica media in cerebral autosomal dominant arteriopathy with subcortical infarcts and leukoencephalopathy. Brain Res, 2012. 1456: p. 64–71.

84. Omar, R., F. Malfait, and T. Van Agtmael, Four decades in the making: Collagen III and mechanisms of vascular Ehlers Danlos Syndrome. Matrix Biol Plus, 2021. 12: p. 100090.

85. Kukita, K., et al., Type IV collagen expression is regulated by Notch3-mediated Notch signaling during angiogenesis. Biochemical and Biophysical Research Communications, 2025. 749: p. 151351.

86. Tefft, J.B., et al., Notch1 and Notch3 coordinate for pericyte-induced stabilization of vasculature. Am J Physiol Cell Physiol, 2022. 322(2): p. C185–c196.

87. Wu, H. and J.T. Parsons, Cortactin, an 80/85-kilodalton pp60src substrate, is a filamentous actin-binding protein enriched in the cell cortex. J Cell Biol, 1993. 120(6): p. 1417–26.

88. Lian, Z., et al., Notch3 enhances the synergistic effect of all-trans retinoic acid and calcipotriol in pancreatic stellate cell activation. J Transl Med, 2025. 23(1): p. 694.

89. Zhao, X., et al., Selective vulnerability of cerebral vasculature to NOTCH3 variants in small vessel disease and rescue by phosphodiesterase-5 inhibitor. Science Advances, 2026. 12(14): p. eaeb1134.

90. Martens, J.H., et al., Cascade of distinct histone modifications during collagenase gene activation. Mol Cell Biol, 2003. 23(5): p. 1808–16.

91. Lee, C., et al., Vascular endothelial growth factor signaling in health and disease: from molecular mechanisms to therapeutic perspectives. Signal Transduction and Targeted Therapy, 2025. 10(1): p. 170.

92. Guo, M.L., et al., Notch3/VEGF-A axis is involved in TAT-mediated proliferation of pulmonary artery smooth muscle cells: Implications for HIV-associated PAH. Cell Death Discov, 2018. 4: p. 22.

93. Lin, S., et al., Non-canonical NOTCH3 signalling limits tumour angiogenesis. Nat Commun, 2017. 8: p. 16074.

94. Ragot, H., et al., Loss of Notch3 Signaling in Vascular Smooth Muscle Cells Promotes Severe Heart Failure Upon Hypertension. Hypertension, 2016. 68(2): p. 392–400.

95. Sansone, P., et al., Self-renewal of CD133(hi) cells by IL6/Notch3 signalling regulates endocrine resistance in metastatic breast cancer. Nat Commun, 2016. 7: p. 10442.

96. Jung, J.G., et al., Notch3 interactome analysis identified WWP2 as a negative regulator of Notch3 signaling in ovarian cancer. PLoS Genet, 2014. 10(10): p. e1004751.

97. Sun, X., et al., NOTCH3 promotes docetaxel resistance of prostate cancer cells through regulating TUBB3 and MAPK signaling pathway. Cancer Sci, 2024. 115(2): p. 412–426.

98. Lee, S.J., X. Zhang, and M.M. Wang, Vascular accumulation of the small leucine-rich proteoglycan decorin in CADASIL. Neuroreport, 2014. 25(13): p. 1059–63.

99. Ishiko, A., et al., Notch3 ectodomain is a major component of granular osmiophilic material (GOM) in CADASIL. Acta Neuropathol, 2006. 112(3): p. 333–9.

100. Peters, N., et al., CADASIL-associated Notch3 mutations have differential effects both on ligand binding and ligand-induced Notch3 receptor signaling through RBP-Jk. Exp Cell Res, 2004. 299(2): p. 454–64.

101. Takahashi, K., et al., Mutations in NOTCH3 cause the formation and retention of aggregates in the endoplasmic reticulum, leading to impaired cell proliferation. Human Molecular Genetics, 2010. 19(1): p. 79–89.

102. Marx, S.O., H. Totary-Jain, and A.R. Marks, Vascular Smooth Muscle Cell Proliferation in Restenosis. Circulation: Cardiovascular Interventions, 2011. 4(1): p. 104–111.

103. Moses, J.W., et al., Sirolimus-eluting stents versus standard stents in patients with stenosis in a native coronary artery. N Engl J Med, 2003. 349(14): p. 1315–23.

104. Marx, S.O., et al., Rapamycin-FKBP inhibits cell cycle regulators of proliferation in vascular smooth muscle cells. Circulation research, 1995. 76(3): p. 412–417.

105. Poon, M., et al., Rapamycin inhibits vascular smooth muscle cell migration. The Journal of clinical investigation, 1996. 98(10): p. 2277–2283.

